# Regulation of Brain Primary Cilia Length by MCH Signaling: Evidence from Pharmacological, Genetic, Optogenetic and Chemogenic Manipulations

**DOI:** 10.1101/2021.04.07.438846

**Authors:** Wedad Alhassen, Yuki Kobayashi, Jessica Su, Brianna Robbins, Henry Ngyuen, Thant Myint, Micah Yu, Surya M. Nauli, Yumiko Saito, Amal Alachkar

**Affiliations:** Departments of Pharmaceutical Sciences, School of Pharmacy, University of California-Irvine, CA 92697, USA; Graduate School of Integrated Sciences for Life, Hiroshima University, 1-7-1 Kagamiyama, Higashi-Hiroshima, Hiroshima, 739-8521, Japan; Department of Biomedical and Pharmaceutical Sciences, School of Pharmacy, Chapman University, Health Science Campus, Chapman University, Irvine, California 92618, USA; Institute for Genomics and Bioinformatics, School of Information and Computer Sciences, University of California-Irvine, CA 92697, USA

**Keywords:** Cilia, melanin concentrating hormone, signaling, brain, activation, inactivation

## Abstract

The melanin concentrating hormone (MCH) system is involved in numerous functions including energy homeostasis, food intake, sleep, stress, mood, aggression, reward, maternal behavior, social behavior, and cognition. MCH acts on a G protein-coupled receptor MCHR1, which expresses ubiquitously in the brain and localizes to neuronal primary cilia. Cilia act as cells’ antennas and play crucial roles in cell signaling to detect and transduce external stimuli to regulate cell differentiation and migration. Cilia are highly dynamic in terms of their length and morphology; however, it is not known if cilia length is causally regulated by MCH system activation in-vivo. In the current work, we examined the effects of the activation and inactivation of MCH system on cilia lengths by using different methodologies, including pharmacological (MCHR1 agonist and antagonist GW803430), germline and conditional genetic deletion of MCHR1 and MCH, optogenetic, and chemogenetic (Designer Receptors Exclusively Activated by Designer Drugs (DREADD)) approaches. We found that stimulation of MCH system either directly through MCHR1 activation, or indirectly through optogenetic and chemogenetic-mediated excitation of MCH neurons, causes cilia shortening. Contrarily, inactivation of MCH signaling through pharmacological MCHR1 blockade or through genetic manipulations - germline deletion of MCHR1 and conditional ablation of MCH neurons - induces cilia lengthening. Our study is the first to uncover the causal effects of the MCH system in the regulation of the length of brain neuronal primary cilia. These findings place MCH system at a unique position in the ciliary signaling in physiological and pathological conditions, and implicate cilia MCHR1 as a potential therapeutic target for the treatment of pathological conditions characterized by impaired cilia function.

## 1. Introduction

Melanin concentrating hormone (MCH), a 19 amino acid hypothalamic neuropeptide, is involved in numerous functions including food intake, energy homeostasis, arousal, sleep, learning and memory, cognition, emotions, and maternal behavior [1–12]. Dysregulation of MCH system has been linked to psychiatric and neurological disorders such as depression and anxiety, and most recently schizophrenia and Alzheimer’s disease [13, 14].

MCH exerts its actions through binding and activating the G-protein coupled receptor (GPCR) MCHR1 that is widely distributed throughout the brain [15, 16]. MCHR1 couples to Gi/o and Gq, resulting in the activation of signaling pathways including Ca2+ mobilization, ERK phosphorylation, and inhibition of cyclic AMP generation. MCHR1 has been shown recently to be selectively located on neuronal primary cilia [17–19], protrusions from the cell bodies of neurons that act as antennas for the cells. Cilia are responsible for signaling functions including detecting and transducing external stimuli that are crucial for development and maintenance of homeostasis (see [20, 21] for reviews). Despite the limited understanding of primary cilia functions, their abundant distribution throughout the brain and the correlation of cilia dysfunctions to some cognitive and behavioral abnormalities and diseases suggest an important role of cilia in brain functions [6]. A few G-protein coupled receptors (GPCRs) have been shown to selectively localize to cilia [22]. We recently showed that dysregulations of cilia genes including ciliary GPCRs are associated with major psychiatric disorders including schizophrenia, autism, major depressive disorder and bipolar disorder [23]. Therefore, MCHR1 localization on cilia membranes might be the basis of the uniqueness of these receptors in the regulating ciliary signaling. In support of this notion, in vitro studies showed that MCH treatment shortened the length of cilia in human retinal pigmented epithelial (hRPE1) cells that were transfected with MCHR1, and that the reduction in cilia length was mediated via Gi/o-Akt pathway [24]. A recent study by Saito group showed that treatment of hippocampus slice cultures with MCH induced cilia shortening in the CA1 region, and that in fasting mice, a marked increase in MCH mRNA expression in the lateral hypothalamus was correlated with a reduction of MCHR1-positive cilia lengths in the hippocampal CA1 region [17]. Therefore, we aimed at establishing the causal relationship between MCH system signaling and cilia length. We examined how the manipulations of the MCH system affect cilia integrity in multiples mouse brain regions. We used multiple techniques to manipulate the MCH system including pharmacological, genetic, optogenetic and chemogenetic approaches to determine how alterations in the MCH system affect cilia length.

## 2. Material and Methods

### 2.1. Ex-vivo study

#### Rat brain slice culture from striatum (caudate/putamen (CPu)) and prefrontal cortex (PFC)

For rat brain slices, Wistar rats (Charles River Japan, Yokohama, Japan) were maintained in a room under a 12 h light:12 h darkness cycle and controlled temperature (23 to 25°C), with water and food available ad libitum. All experimental protocols were reviewed by Hiroshima University Animal Care Committee and met the Japanese Experimental Animal Research Association standards, as defined in the Guidelines for Animal Experiments (1987). Newborn rats were anesthetized and decapitated for slice culture preparation. We established rat CPu and PFC slice culture method for clear detection of primary cilia and ciliary MCHR1 based on a previous study [17]. The brains of 11-day-old rats were rapidly removed, and the hippocampi were dissected and cut into 200-µm coronal slices using a McIlwain tissue chopper (The Mickle Laboratory Engineering, Surrey, UK). The slices were trimmed for tissue in ice-cold dissection buffer containing 2.5 mM KCl, 0.05 µM CaCl2, 1.7 mM NaH2PO4, 11 mM glucose, 20 mM HEPES, 8 mM NaOH, 18 mM NaCl, 0.23 M sucrose, and PG/SM, and placed on 0.4-µm, 30- mm diameter semiporous Millicell Cell Culture inserts (PICM0RG50; Merck Millipore, Germany) in 35-mm petri dishes. Each dish contained 1 ml of starter medium comprising 50% MEM (GIBCO, Grand Island, NY, USA), 25% normal horse serum, 25% Hanks’ Balanced Salt Solution, 36 mM glucose, and PG/SM. Six hippocampal slices were placed on each membrane, and maintained at 37°C in a 5% CO_2_ incubator. After 1 day, the medium was changed to Neurobasal Medium (21103-049; Gibco) containing 1.2% B27 Serum-Free Supplement (17504- 044; Gibco), 25 mM GlutaMAX I (Gibco), and penicillin G sodium/streptomycin sulfate (Slice- GM). One-third of the Slice-GM was changed every 72 h. The CPu and PFC slices were treated with MCH (30 nM, Peptide Institute, Osaka, Japan) in Slice-GM and fixed on day 14 or 7 of culture, respectively. Immunohistochemical staining of hippocampal slices was performed in 6- well culture plates. The slices were fixed for 3 h at 4°C in fresh 4% paraformaldehyde, washed with D-PBS, heated (70°C for 20 min) in 10% Histo VT One (Nacalai Tesque, Kyoto, Japan) in D-PBS for antigen retrieval, and blocked with D-PBS containing 5% horse serum and 0.1% Triton X-100 for 2 h. The slices were then incubated with rabbit anti-rat adenylate cyclase 3 (ADCY3, RPCA-ACIII; Encor Biotechnology, Alachua, FL, USA; 1:5000) and goat anti-human MCHR1 (goat anti-human MCHR1; C-17, sc-5534; Santa Cruz Biotechnology; 1:300) primary antibodies overnight at 4°C. The bound antibodies were detected by incubation with Alexa Fluor 488-conjugated donkey anti-rabbit IgG (Thermo Fisher Scientific, Rockford, IL, USA; 1:300) or Alexa Fluor 546-conjugated donkey anti-goat IgG (Thermo Fisher Scientific; 1:300) secondary antibodies for 2 h at room temperature. The slices were counterstained with DAPI for 10 min and mounted with VECTOR Shield.

#### Microscopic images and analysis

The primary cilia lengths was measured with a BZ-9000 fluorescence microscope (Keyence, Osaka, Japan) using PhotoRuler Ver. 1.1 software (The Genus Inocybe, Hyogo, Japan). A minimum of 80 cilia per treatment were obtained from at least three independent experiments, and the values are presented as means ± SEM. The scatter plot graphs were created by using the box plot and beeswarm module of a statistical software R (https://cran.r-project.org/).

### 2.2. In-vivo Studies

All experimental procedures were approved by the Institutional Animal Care and Use Committee of the University of California, Irvine and were performed in compliance with national and institutional guidelines for the care and use of laboratory animals.

#### 2.2.1. Pharmacological Manipulations

##### Intracerebroventricular administration of MCH

Eight week old mice (n=16) underwent stereotaxic surgery for implantation of a stainless steel guide cannula into the lateral ventricles (20 gauge guide cannulas with 2.5-mm custom-cut depth, PlasticsOne). Animals were anesthetized with 2% isofluorane anesthesia (Institutiional Animal Care and Use Committee guidelines) and were secured in a Kopf steroeotaxic instrument. Guide cannula was implanted at -0.22mm posterior to bregma, 1.0 mm lateral, and 2.3 mm below the skull surface. Post-surgery animals were allowed to recover for one week with dummy cannula in place before injections. Animals were then infused with either vehicle or MCH peptide (1nmol) for 7 consecutive days using a 50 µl Hamilton microsyringe. MCH (1 nmol) was dissolved in phosphate buffered saline (pH7.4) with 0.2% bovine serum albumin. The dose of MCH was determined by previous reported findings [25]. Following the final injection animals were perfused.

##### Intraperitoneal administration of MCHR1 antagonist GW 803430

Eight-week Swiss Webster animals (n=16) were administered intraperitoneally (i.p.) 3mg/kg MCH1R antagonist GW803430 or vehicle for 7 consecutive days [25, 26]. This dose was selected based on previously receptor occupancy studies demonstrating that near complete blockade of the MCH system is achieved following i.p. administration at the 3mg/kg dose [25] [26]. Following the final injection, animals were perfused transcardially for tissue fixation.

#### 2.2.2. Genetic Manipulation

##### Germline MCHR1 Knockout (MCHR1 KO) and MCH Conditional Knockout (MCH cKO) mice

MCHR1 KO mice were generated as previously described [27]. PmchCre/+;iDTR/^+^ mice were generated as described previously [28]. Briefly, Pmch-Cre mice (Jackson Laboratories, Bar Harbor, Maine, USA) that express Cre-recombinase (Cre) under the MCH promoter [29] were crossed with homozygous inducible diphtheria toxin receptor iDTR/^+^ mice (from Dr. Satchidinanda Panda and originally generated in the lab of Dr. Ari Waisman) [13], [25], [28], [30]. The resulting iDTR^+^PmchCre^+^ (iDTR^+^/Cre^+^) and their control littermate iDTR^+^PmchCre^-^ (iDTR^+^/Cre^-^) mice were injected twice in four days with the diphtheria toxin (DT) (16µg/kg, i.p).

#### 2.2.3. Optogenetic stimulation of MCH neurons

##### Surgery

AAV5-EF1a-DIO-ChR2-T159c-eYFP (titer: ≥ 1×10¹³ vg/mL) is an EF1a-driven, Cre-dependent, humanized channelrhodopsin E123T/T159C mutant fused to EYFP for optogenetic activation (pAAV-Ef1a-DIO hChR2 (E123T/T159C)-EYFP was a gift from Karl Deisseroth (Addgene viral prep # 35509-AAV5; http://n2t.net/addgene:35509; RRID: Addgene_35509) [31]. AAV5-EF1a- DIO-ChR2-T159c-eYFP was stereotaxically injected into the lateral hypothalamus (flat skull coordinates from bregma: anteroposterior, -1.30mm, mediolateral, +1mm, and dorsalventral, - 5.20mm) of 8-week-old transgenic C57BL/6 PmchCre + (n=8) and PmchCre - animals (n=8), (Jackson Laboratories, Bar Harbor, Maine, USA) that express Cre-recombinase (Cre) under the MCH promoter. One injection in each mouse (one hemisphere; 800 nl). At this time the high-power LED fiber cannula (core diameter 200 µm, outer diameter 225 µm, length 2.5 mm, numerical aperture 0.66, Prizmatix, Israel) was also implanted in the lateral hypothalamus at - 5.0mm slightly above injection site. The cannula light irradiance was adjusted to 6 mW before implantation. An additional hole was drilled on the opposite side for stainless steel holding screws. The fiber was fixed with dental cement holding the fiber and screw to the skull. The skin was closed with silk sutures. Animals were allowed to recover for one week before the start of the experiments. Correct fiber placement and injection site was ascertained post mortem on coronal brain sections. Animals were exposed to blue light at 10 ms pulse 1s repeated every 4s for thirty minutes before perfusions after the feeding behavior test.

##### Optogenetic stimulations and behavioral analysis

Animals were habituated to the connection of the fiber patch cords to their cannulas inside their home cage for 1 h for 3 days. On the fourth day animals were connected to the fiber patch cord and were tested in their home cage on feeding behavior. Animals were exposed to 460 nm blue led light source to excite ChR2 at a specific stimulation paradigm: 10 Hz 10ms pulse 1s repeated every 4s for 10 minutes. Feeding behavior was tracked and recorded (Fig S1).

#### 2.2.4. Designer Receptors Exclusively Activated by Designer Drugs (DREADD)-based chemogenetic stimulation of MCH neurons

AAV8-hSyn-DIO-hM3D(Gq)-mCherry (titer:4.8×10^12^ GC/mL) (pAAV-hSyn-DIO-hM3D(Gq)- mCherry was a gift from Bryan Roth (Addgene viral prep #44361-AAV8; http://n2t.net/addgene: 44361; RRID:Addgene_44361)) was stereotaxically injected into the lateral hypothalamus (flat skull coordinates from bregma: anteroposterior, -1.30mm, mediolateral, +1mm, and dorsalventral, -5.20mm) of 8 week old transgenic C57BL/6 PmchCre+ (n=8) and PmchCre- animals (n=8). This virus is a Syn-driven, Cre-dependent, hM3D(Gq) receptor with an mCherry reporter for CNO-induced neuronal activation [32]. Two injections per mouse (both hemispheres; 0.6 nl) were administered. Animals were allowed to recover for one week before the start of the experiments. CNO (1 mg/kg) was administered 90 minutes before perfusions (Fig. 4a).

**Fig.1.**
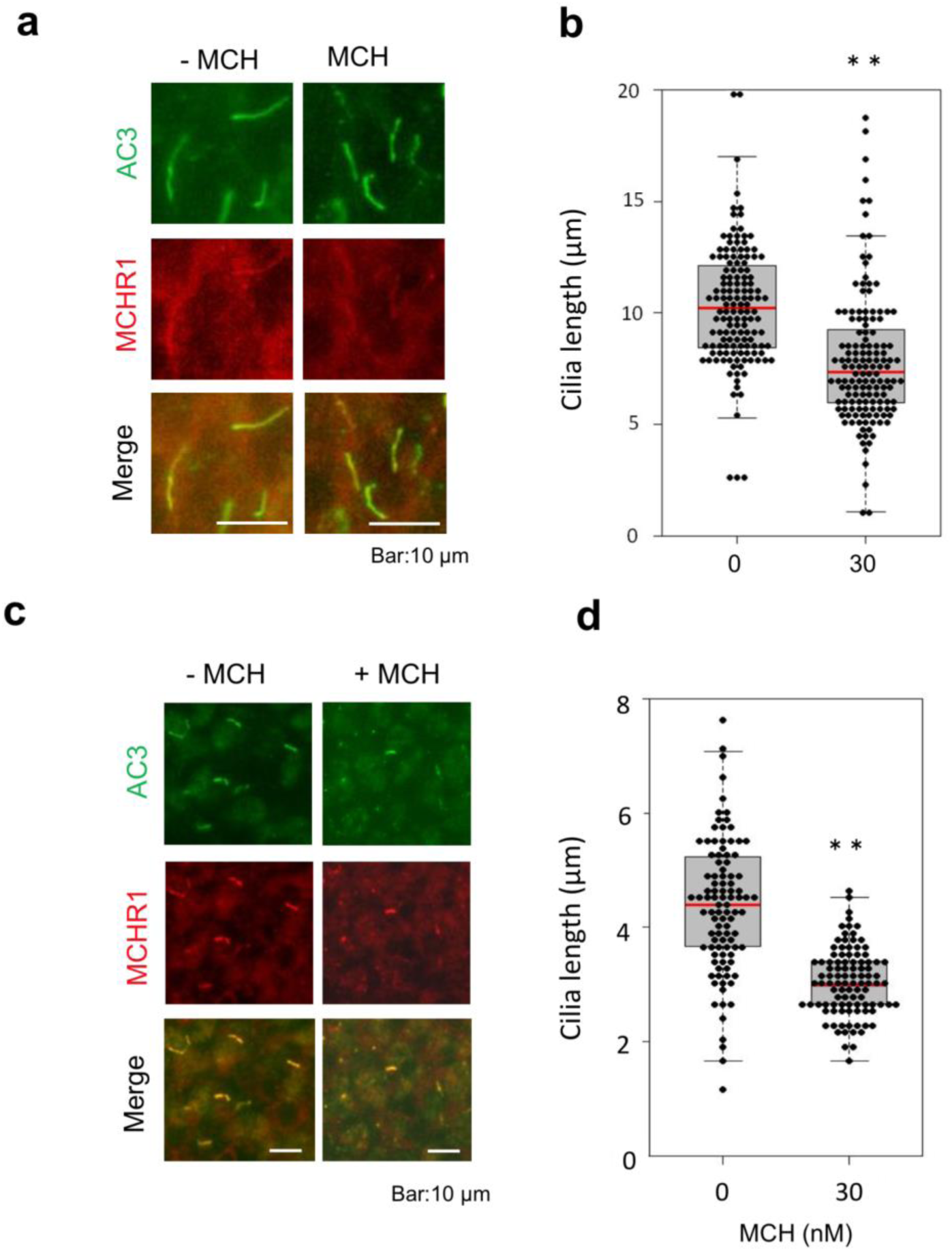
MCH treatment leads to cilia length shortening in cultured brain slices. (a, b) Rat CPu slice cultures were treated with or without 30 nM MCH for 18 h. (c, d) Rat PFC slice cultures were treated with or without 30 nM MCH for 6 h. (a, c) Primary cilia were co- labeled with antibodies against ADCY3 (green) and MCHR1 (red). Scale bars, 10 µm. (b, d) The scatter-plot represents cilium lengths measured in randomly selected fields; at least 80 cilia per group in both CPu slices (b) and PFC slices (d) were evaluated, respectively. Primary cilia were significantly shorter in MCH treated cultures than in control cultures for both CPu slices and PFC slices. \*\**P* < 0.01.

#### 2.2.5. Immunohistochemistry

Immunofluorescent staining was carried out as previously reported [33]. Briefly, animals were perfused transcardially under isofluorane anesthesia with saline followed by 4% paraformaldehyde in phosphate buffer saline brains were removed and coronal brain sections (30 µm) were cut. The MCH-neuron ablation was verified by visualizing MCH neurons using rabbit polyclonal anti-MCH antibody (1:150,000, antibody courtesy of W. Vale, Salk Institute, La Jolla, CA, USA) as previously described [25]. A goat anti-rabbit (1:500; ThermoFisher) was used to visualize MCH immunoreactivity. For cilia study, three to five sections were selected from each region of interest according to the mouse brain atlas [20]. Primary cilia were stained with ADCY3 (rabbit, 1:500; Santa Cruz Biotechnology) and the secondary antibody donkey anti-rabbit Alexa 546 (Thermo Fisher Scientific). For c-Fos immunohistochemistry, mice were perfused 90 minutes after stimulation and brain sections were stained with rabbit anti-c-Fos antibody (1:500, Invitrogen). Nuclei were stained with 4’,6-diamidino-2-phenylindole (DAPI) solution (1:10000), and mounted with Aquamount mounting solution. Image acquisition was carried out using confocal laser microscope. Images were captured using Leica Sp8 TCS confocal microscope (UCI optical biology core facility). Cilia length was measured using ImageJ [34] in all cells in each section, and the mean values of three sections per brain of 3-5 brains were calculated. All cilia measurements were performed by two persons blind to the experiment conditions.

### 2.3. Statistical analysis

GraphPad Prism (GraphPad Software, Inc.) was used for statistical analysis, and all data were presented as mean ± standard error mean (SEM). Student’s unpaired t-test was used to analyze the results. *P* value < 0.05 was considered statistically significant.

## 3. Results

### 1. MCH shortens cilia length in the rat striatum and prefrontal cortex culture

Immunohistochemical analyses in brains of rats showed that ADCY3/MCHR1 double-positive neuronal primary cilia were localized in discrete regions including CPu and cerebral cortex [35]. Cilia MCHR1 merge with ADCY3 was also observed in cultured slices derived from the rat CPu and PFC (Fig.1 a,c). The neuronal cilia length in the CPu slices was 10.30 ± 0.23 µm (mean ± SEM), and MCH treatment for 18 h decreased this length by 23.0% (7.90 ± 0.28 µm, Fig. 1b). We observed a similar effect of MCH in rat PFC slice cultures. Exposure to MCH led to neuronal cilia shortening by 31% (4.39 ± 0.12 µm vs 3.05 ± 0.06 µm; mean ± SEM, *P* < 0.01, Fig.1d).

### 2. Activation of MCH system shortens cilia length in the mouse brain

#### 2.1. MCHR1 agonist shortens cilia in the mouse brain

The central (i.c.v) administration of MCH in adult mice for 7 consecutive days caused a significant decrease in the cilia length in several regions of the brain including the hippocampus, striatum, prefrontal cortex (PFC), and nucleus accumbens (NAc). In the PFC, administration of MCH caused a 42% reduction in cilia length (2.85 ± 0.1 µm compared to 4.96 ± 0.30 µm in the PFC of the animals administered vehicle, t=6.433, P=0.0007, unpaired t-test, Fig 2b,c). In the CA1, administration of MCH caused a 33% shortening in cilia length (6.743 ± 0.23 µm compared to 10.18 ± 0.30 µm in the CA1 of the animals administered vehicle, t=8.436, P=0.0002, unpaired t-test, Fig 2d,e). In the CPu, administration of MCH caused a 37% reduction in cilia length (5.04 ± 0.17 µm compared to 8.005 ± 0.2 µm in the CPu of the animals administered vehicle, t=10.03, P<0.0001, unpaired t-test, Fig 2f,g). In the NAc, administration of MCH caused a reduction in cilia length by 32% (5.566 ± 0.33 µm compared to 8.287 ± 0.67 µm in the NAc of the animals administered vehicle, t=3.648, P=0.0107, unpaired t-test, Fig 2h,i). Number of cells (%) were grouped by cilia length (µm) in the PFC, CA1, CPu, and NAc to further show the decrease in cilia length when administered MCH i.c.v (Fig. 2j). Administration of MCH i.c.v. resulted in a significant decrease in cilia density per field in the PFC (t=3.849, P=0.0085), CA1 (t=3.141, P=0.02), CPu (t=4.224, P=0.0055), and NAc (t=3.766, P=0.0093, Fig. S2).

**Fig.2.**
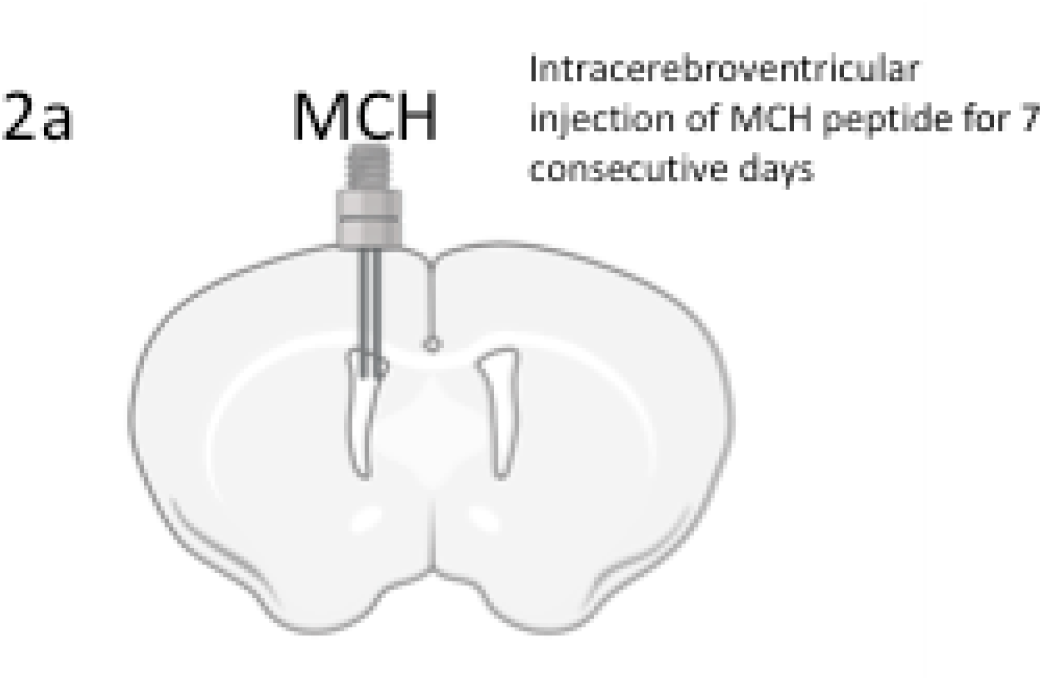
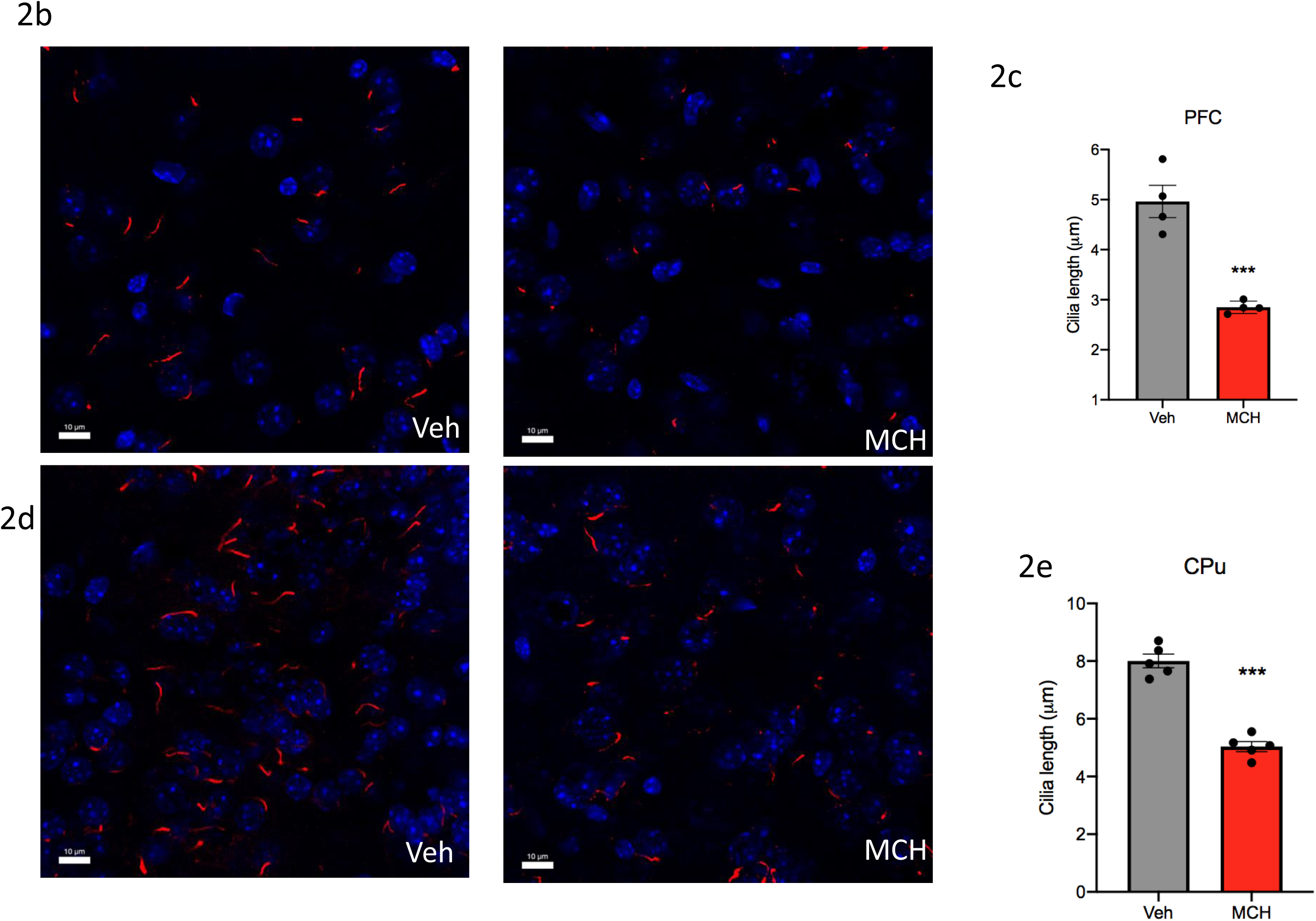
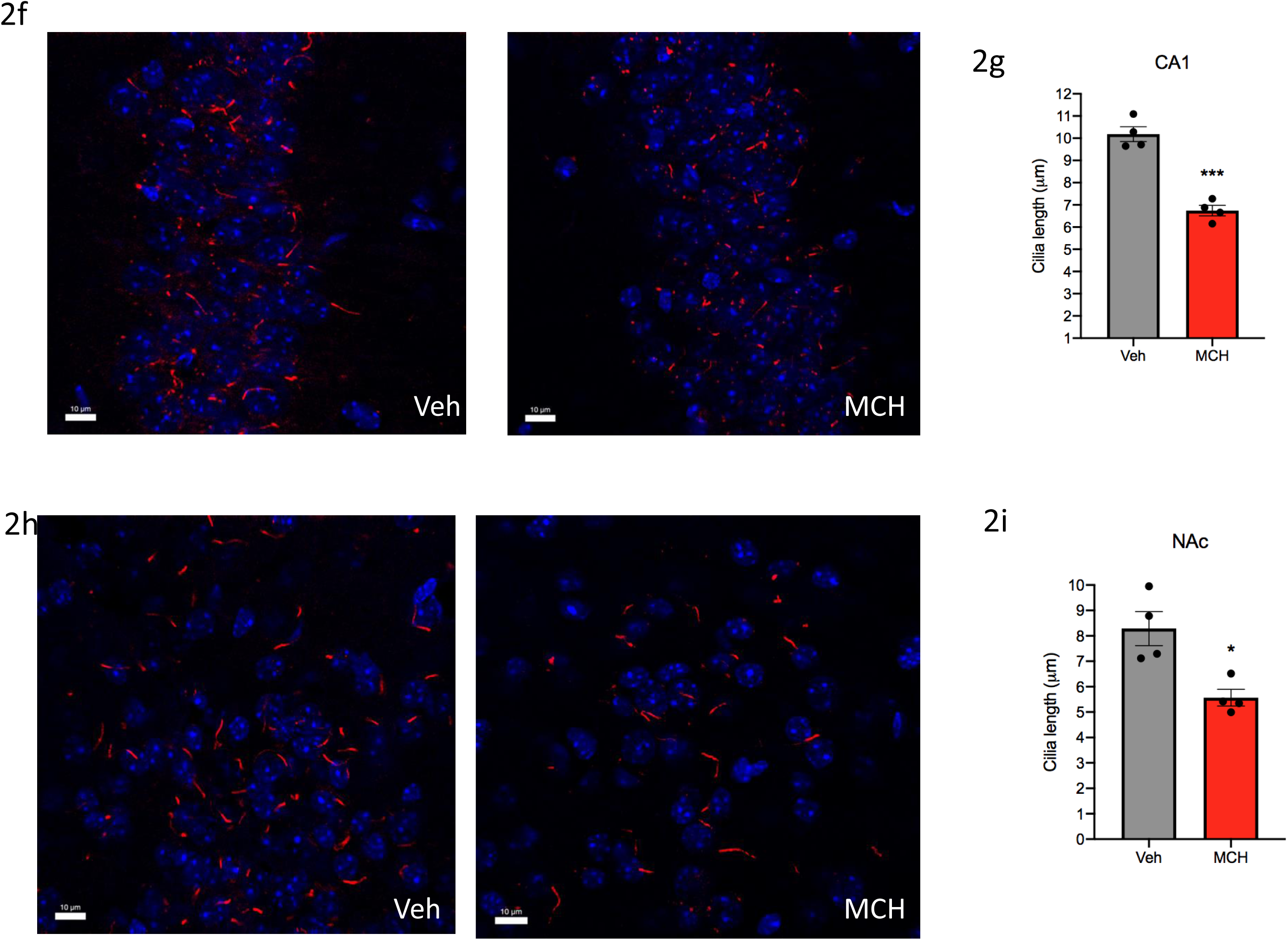
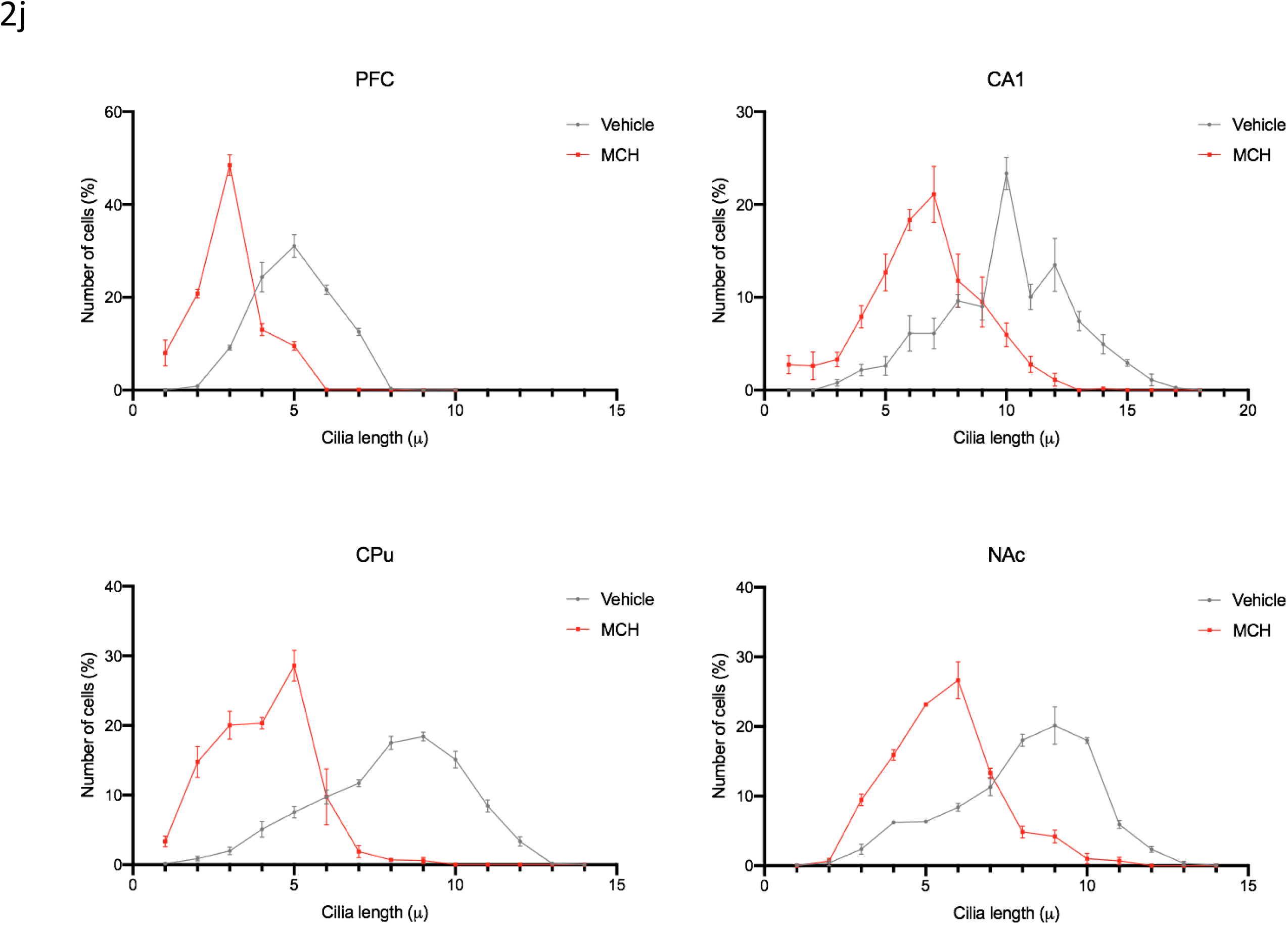
Administration of MCH to mice shortens cilia length. (a) Schematic representation of intracerebroventricular infusions of MCH peptide or vehicle for 7 consecutive days in adult mice. Immunostaining of ADCY3 labeled in red fluorescence in the (b) PFC, (d) CA1, (f) CPu and (h) NAc in animals administered with MCH and vehicle (Scale bar= 10 µm). Quantification of cilia length (µm) in the (c) PFC, (e) CA1, (g) CPu (CPu), (i) NAc (NAc), (j) Grouped cilia length (µm) by number of cells (%) in the PFC, CA1, CPu, and NAc.

#### 2.2. Optogenetic and Chemogenic excitation of MCH neurons shortens cilia length

Optogenetic stimulation of MCH neurons was applied to one hemisphere of the brain at 460 nm blue led light at 10 Hz 10ms pulse 1s repeated every 4s for 10 minutes, based on previously reported rates, at which that MCH neurons fire [36, 37]. MCH neurons were activated through light stimulation of ChR2 in PmchCre+ adult mice, which caused a significant decrease in cilia length in comparison to PmchCre- adult mice (control) in both hemispheres of the brain in several regions of the brain including the CA1, striatum, PFC, and NAc. Co-expression of GFP (green) in the lateral hypothalamic MCH neurons was confirmed using double immunostaining (red) in PmchCre+ and PmchCre- mice (Fig. 3b). Coexpression of GFP fluorescence (green) and c-Fos immunofluorecense (red) identifies cells recently activated neurons in PmchCre+ and PmchCre- mice (Fig. 3c). In the PFC stimulating MCH neurons caused a reduction in cilia length by 59% (6.497 ± 0.6 µm in PmchCre- mice compared to 2.635 ± 0.23 µm in pMchCre+ mice, t=6.319, P=0.0007, unpaired t-test) (Fig 3d, 3e). In the CA1 stimulating MCH neurons caused a significant reduction in cilia length by 34% (3.823 ± 0.33 µm in pMchCre- mice compared to 2.536 ± 0.14 µm in pMchCre+ mice, t=3.535, P=0.0123, unpaired t-test, Fig 3f,g). In the CPu stimulating MCH neurons caused a significant reduction in cilia length by 56% (6.065 ± 0.45 µm in pMchCre- mice compared to 4.031 ± 0.4 µm in pMchCre+ mice, t=3.520, P=0.0078, unpaired t-test, Fig 3h,i). In the NAc stimulating MCH neurons caused significant reduction in cilia length by 20% (7.442 ± 0.46 µm in pMchCre- mice compared to 5.952 ± 0.34 µ in pMchCre+ mice, t=2.593, P=0.0410, unpaired t-test, Fig 3j,k). Number of cells (%) were grouped by cilia length (µm) in the PFC, CA1, CPu, and NAc to further show the decrease in cilia length when MCH is stimulated via optogenetics (Fig. 3l). pMchCre+ mice also exhibited more feeding behavior than pMchCre- mice when exposed to blue light to stimulate MCH neurons. Over the observed 10 minutes, pMchCre+ mice averaged 74.13 seconds of feeding while pMchCre- mice averaged 7.413 seconds (t=7.027, P<0.001) (Fig. S1). Stimulating MCH neurons via optogenetics caused a significant decrease in cilia density per field in the PFC (t=3.789, P=0.0091), CA1 (t=6.324, P=0.0007), and CPu (t=5.656, P=0.0013, Fig. S3).

**Fig.3.**
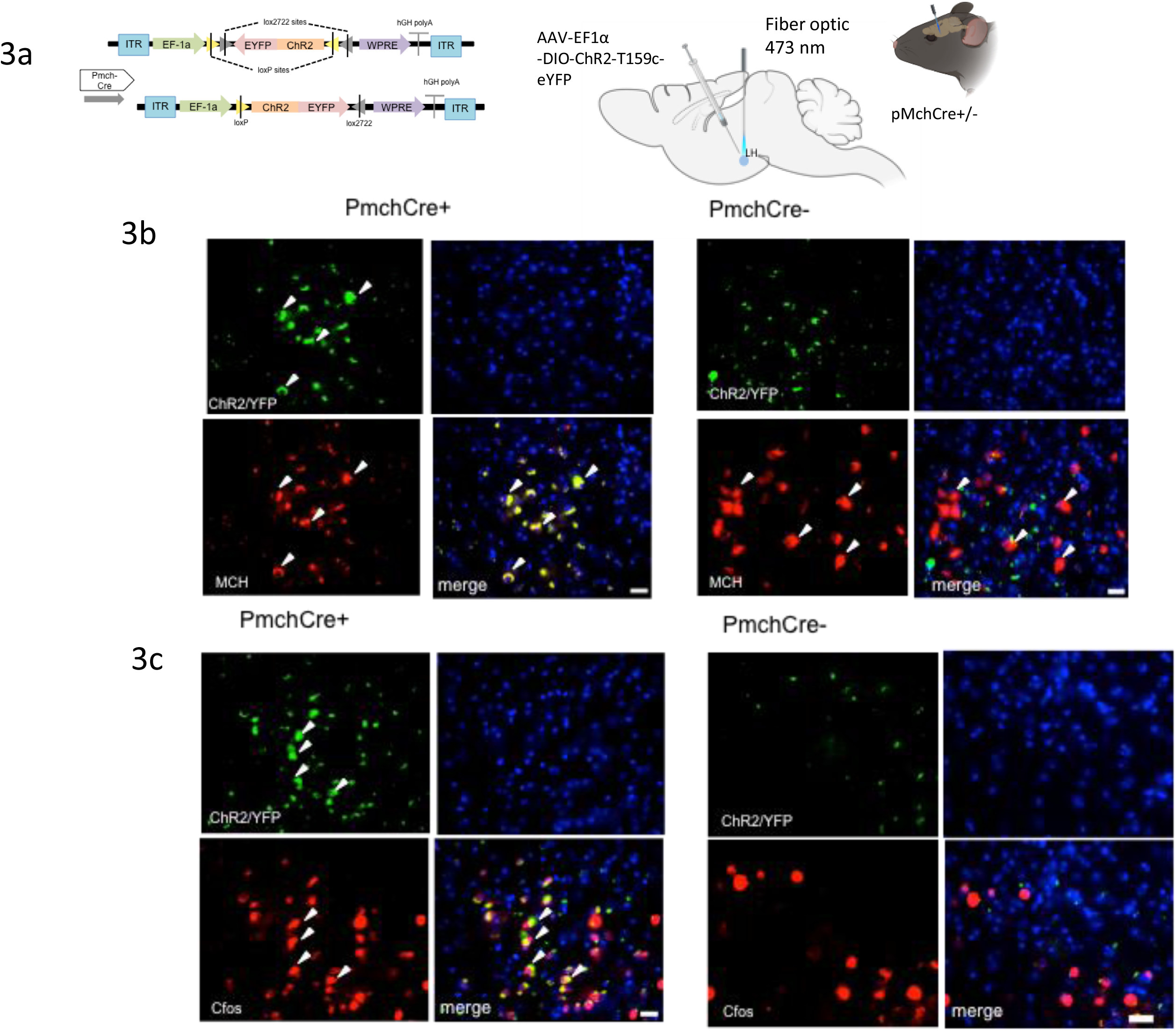
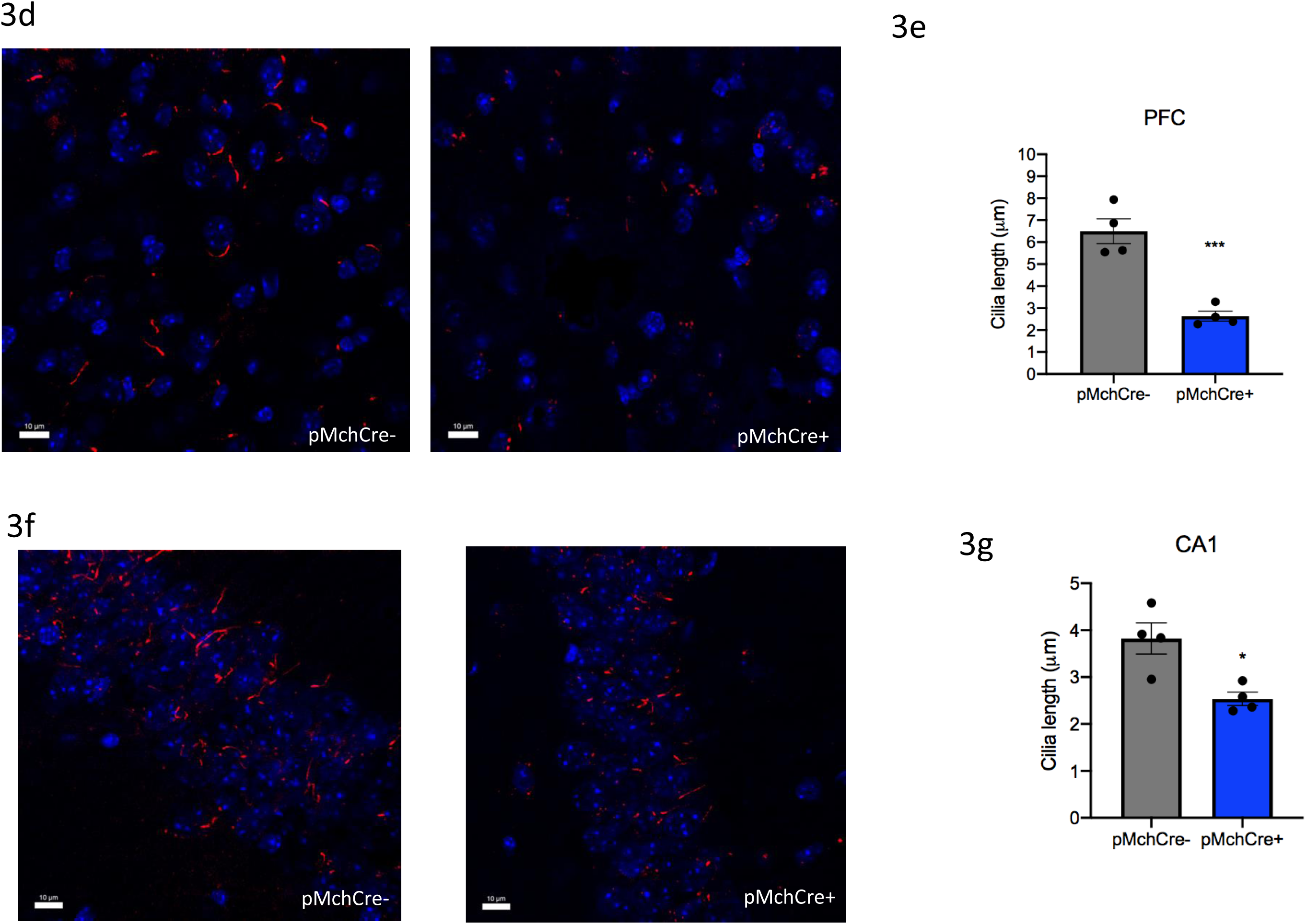
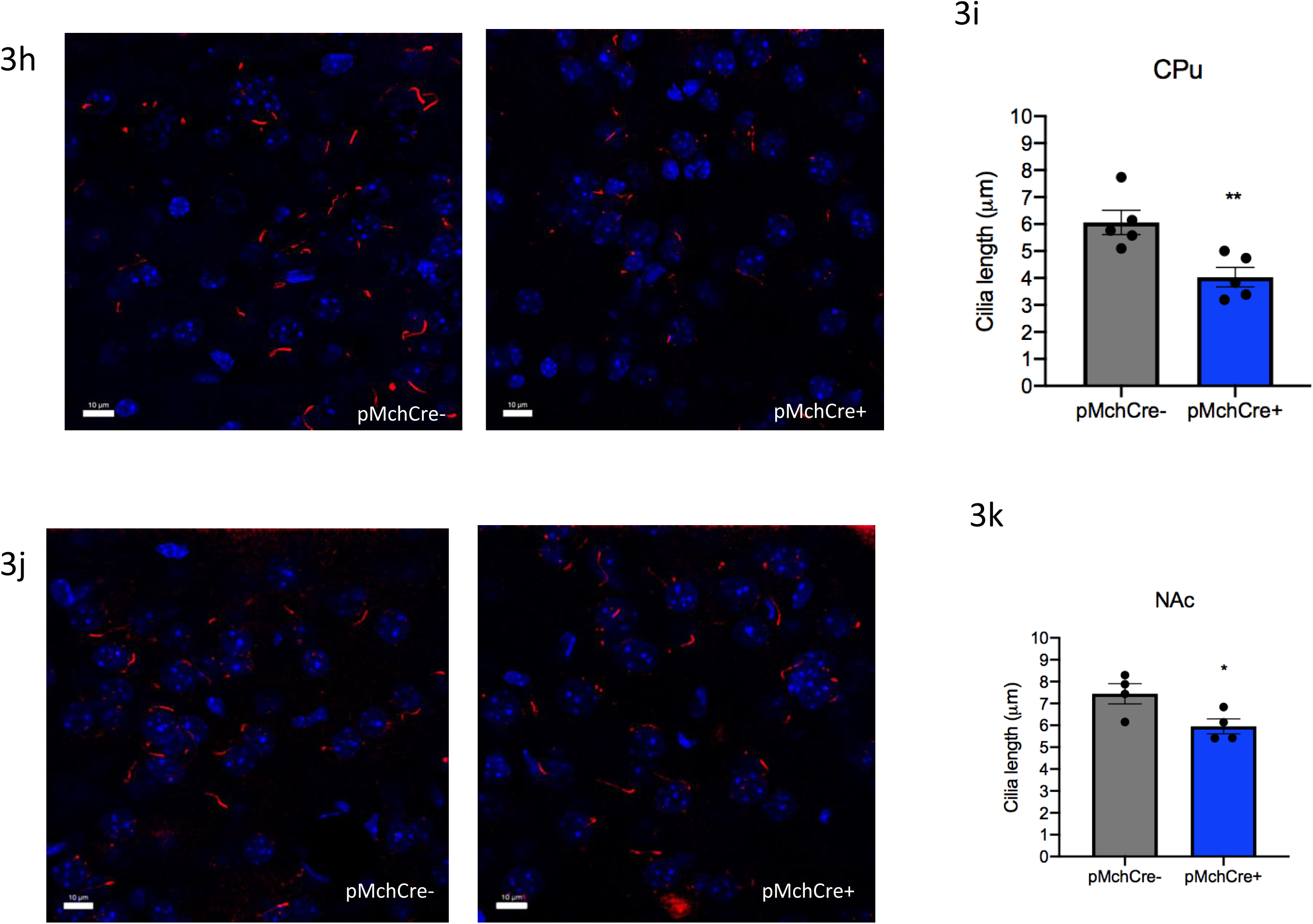
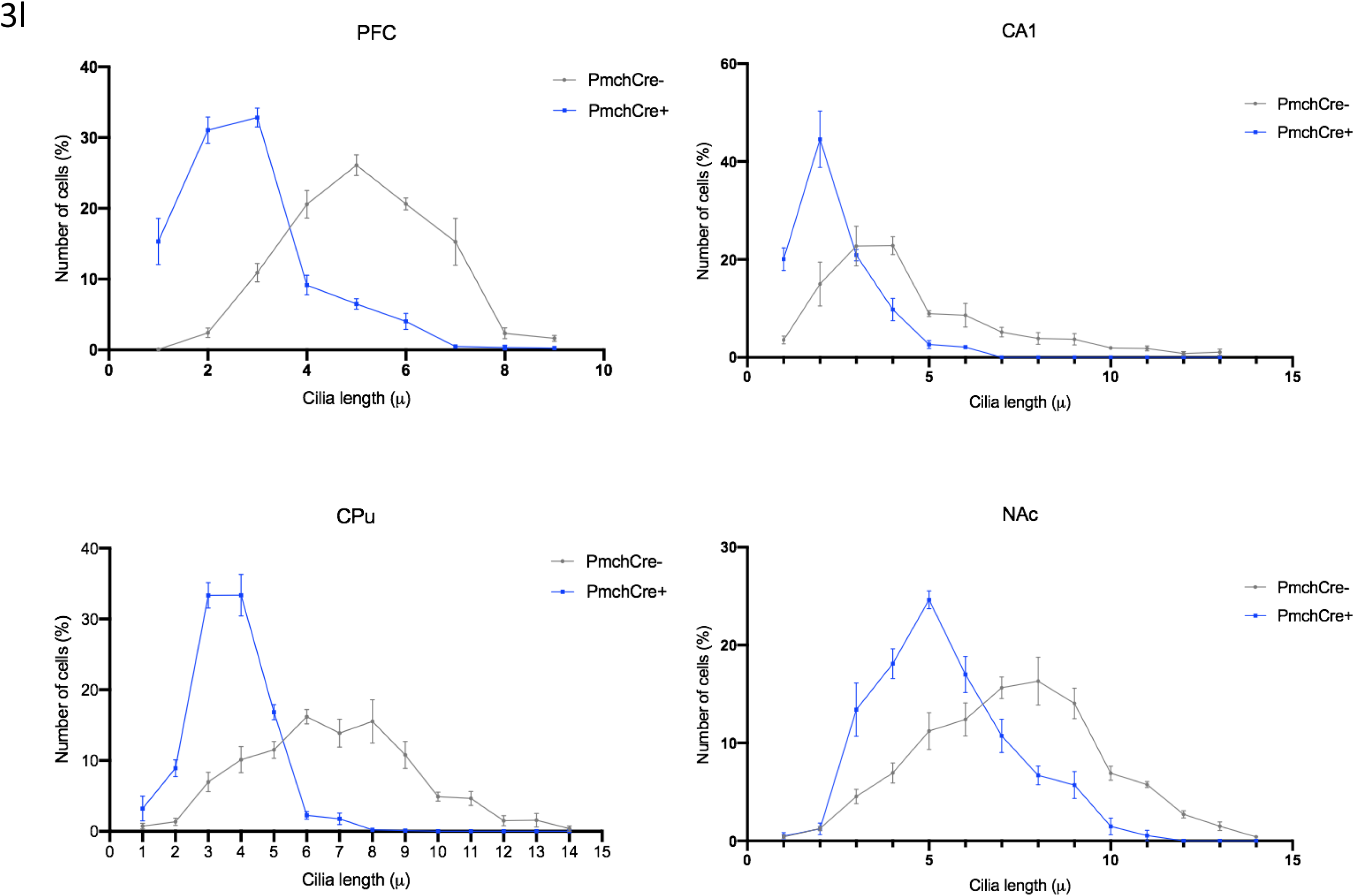
Administration of the GW803430 (MCHR1 antagonist) to mice increases cilia length. (a) Schematic representation of intraperitoneal injection of GW803430 or vehicle for 7 consecutive days in adult mice. Created with BioRender.com Immunostaining of ADCY3 labeled in red fluorescence in the (b) PFC, (d) CA1, (f) CPu and (h) NAc in animals administered with MCH and vehicle (Scale bar= 10 µm). Quantification of cilia length (µm) in the (c) PFC, (e) CA1, (g) striatum (i) NAc. (j) Grouped cilia length (µm) by number of cells (%) in the PFC, CA1, CPu, and NAc.

**Fig. 4.**
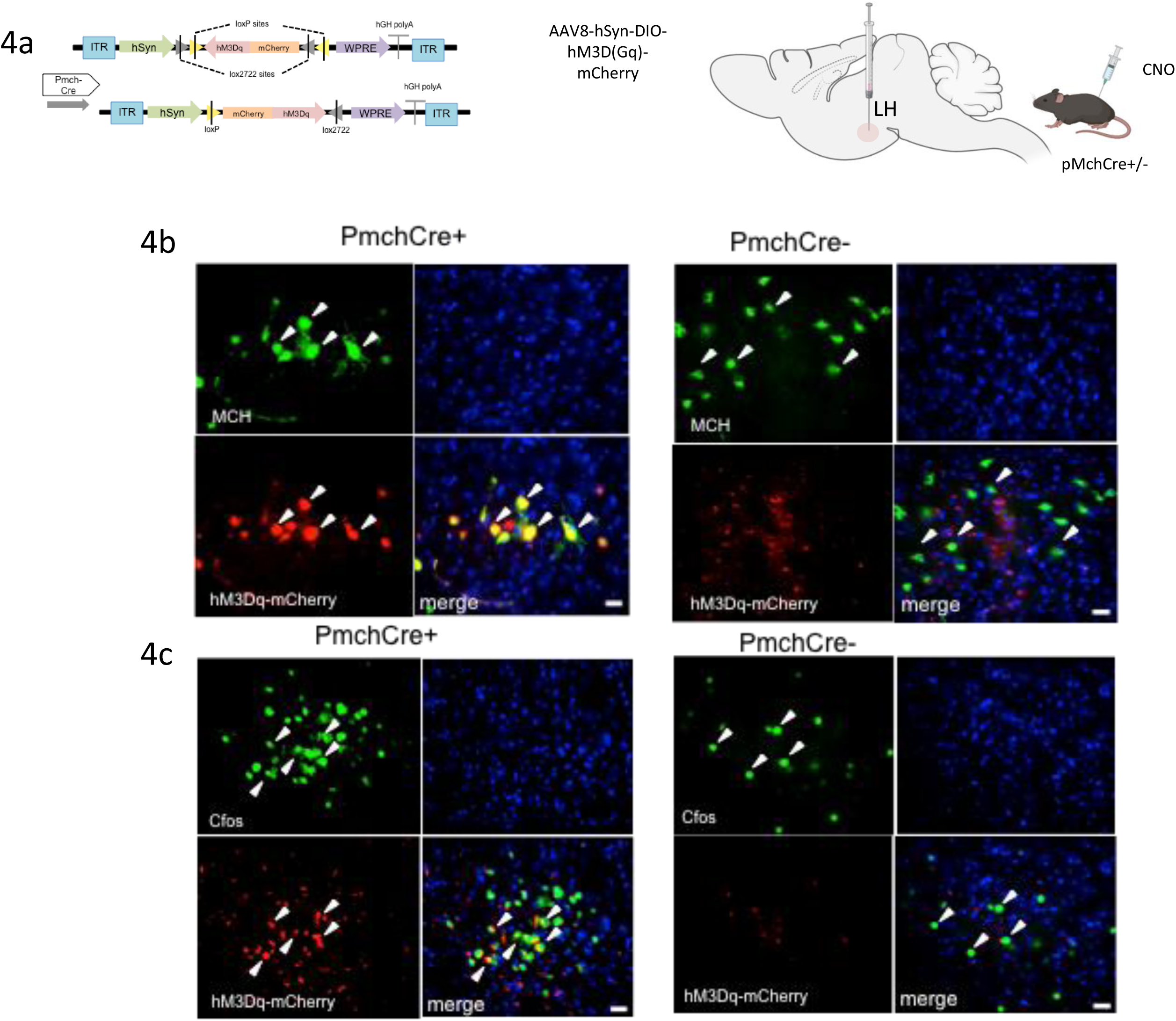
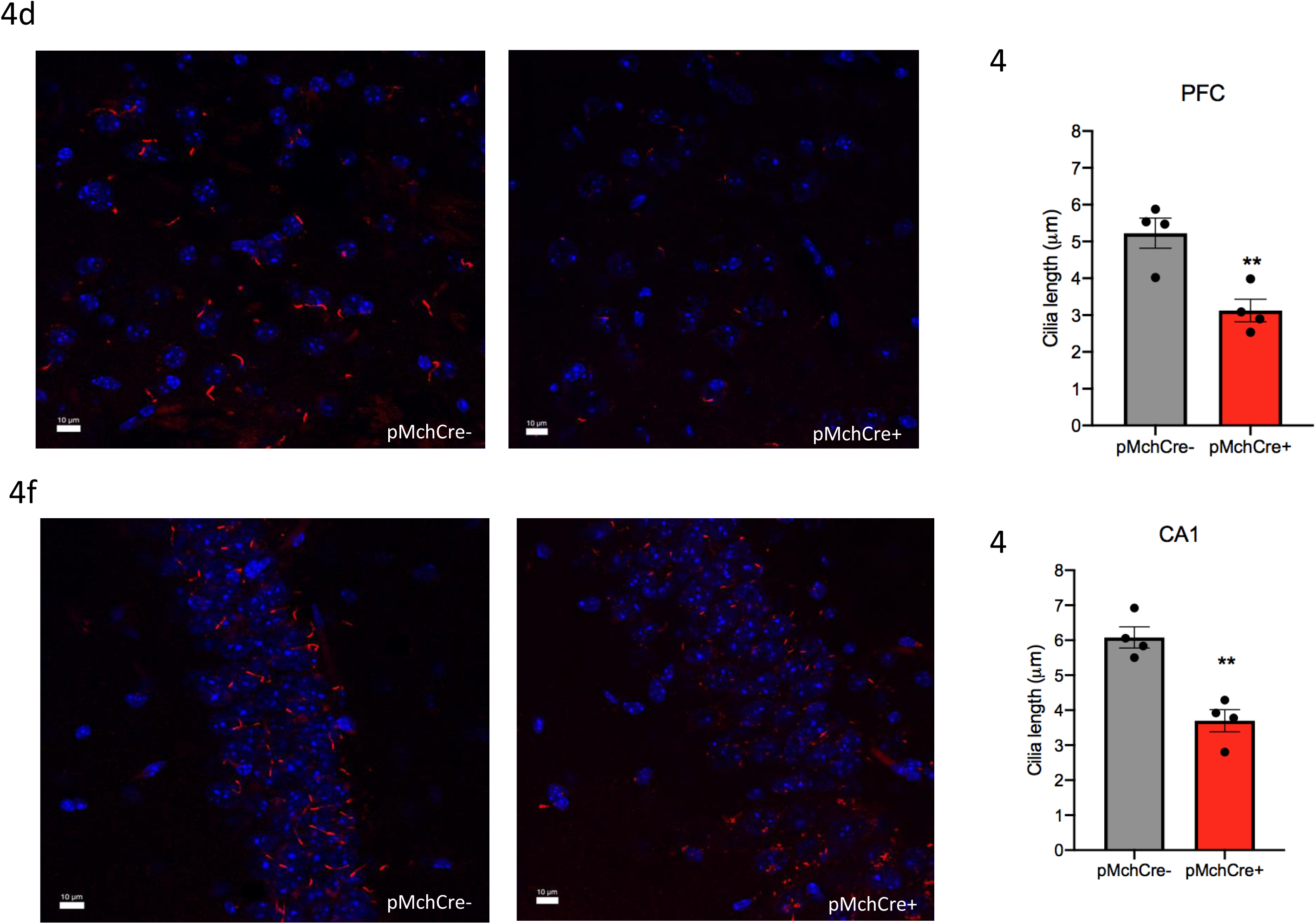
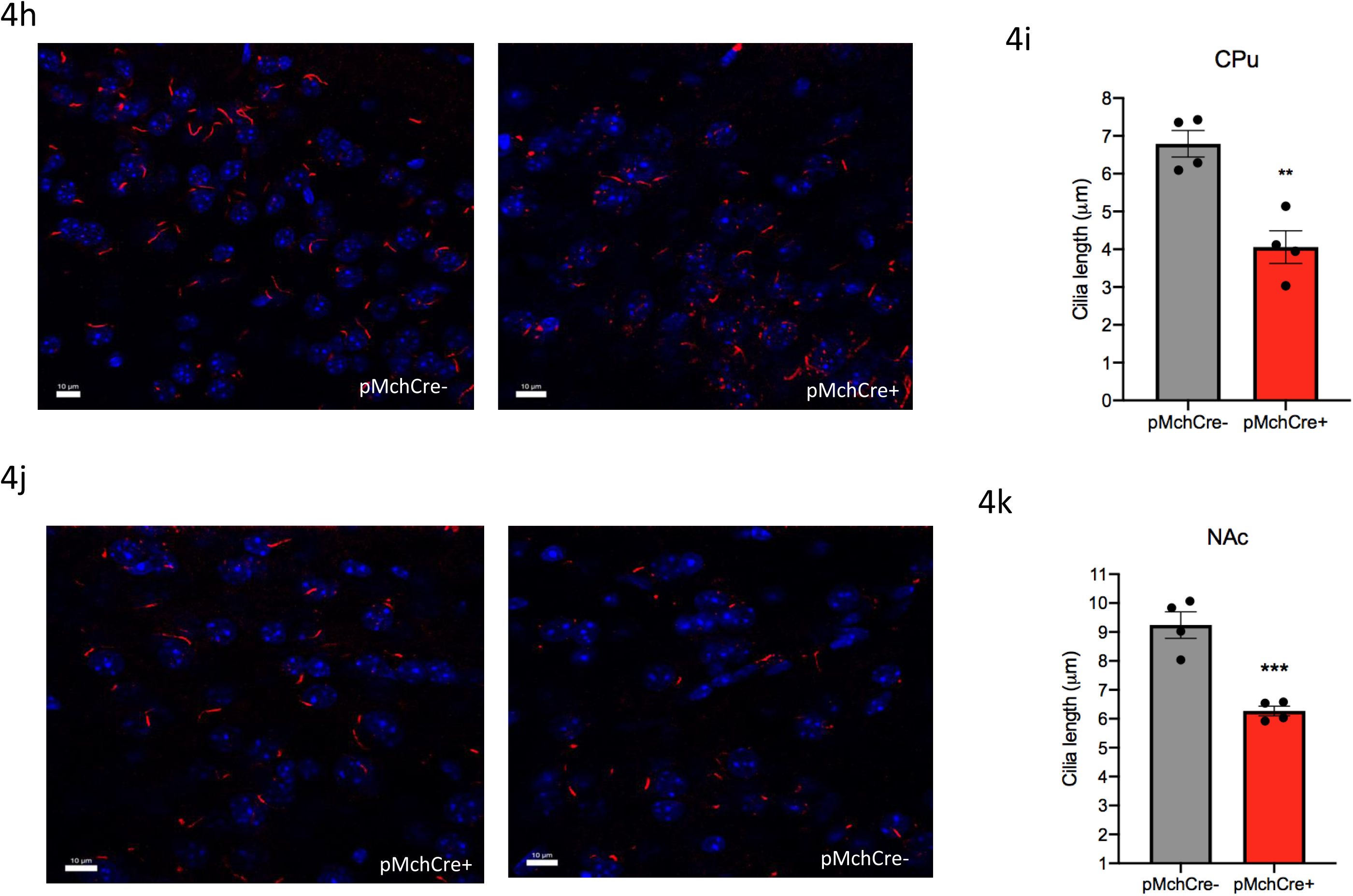
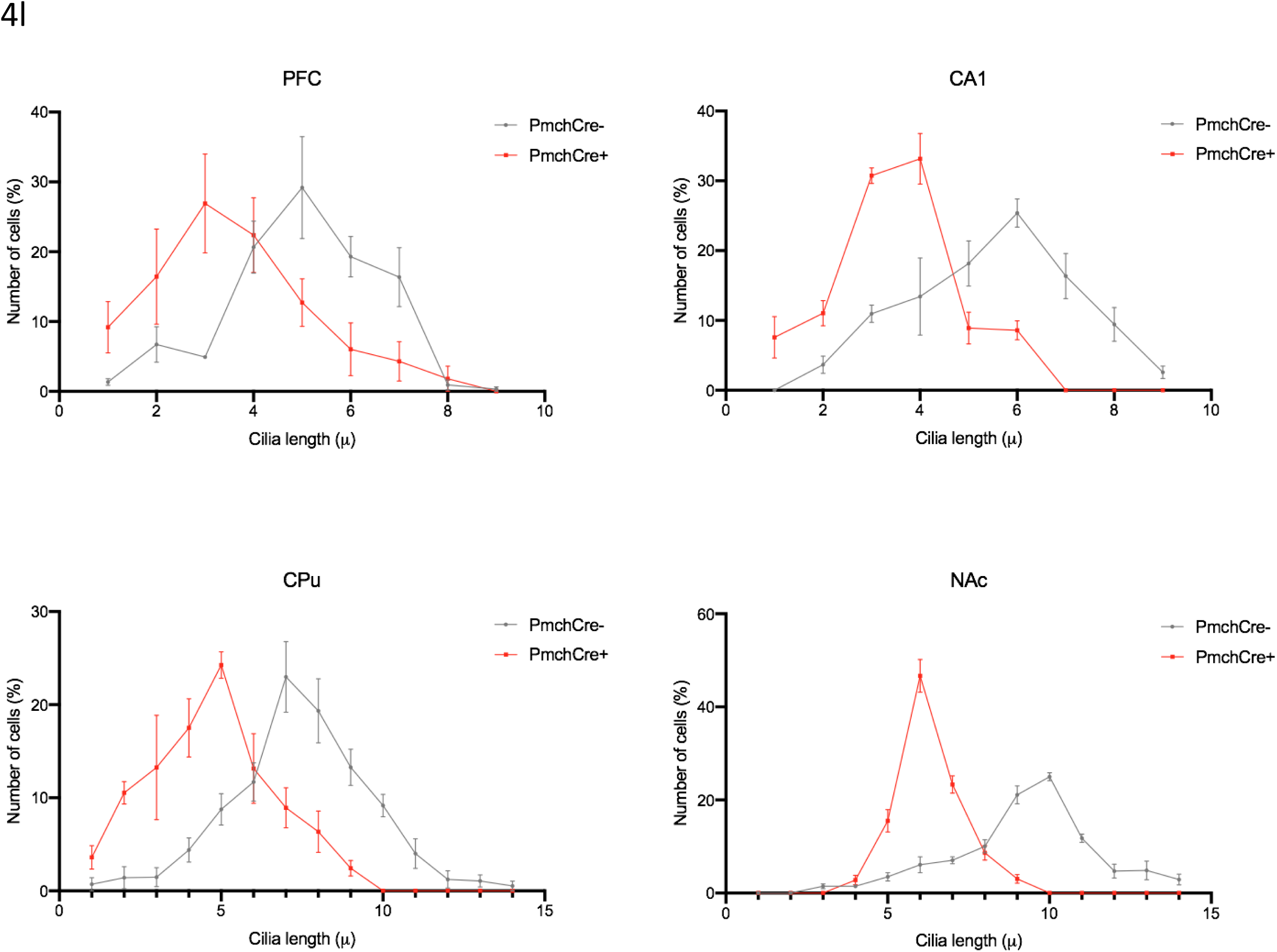
Cilia length is increased in MCHR1 Knockout animals. Immunostaining of ADCY3 labeled in red fluorescence in the (a) PFC, (c) CA1, (e) CPu and (g) NAc in MCHR1 knockout animals (Scale bar= 10 µm). Quantification of cilia length (µm) in the (b) PFC, (d) CA1, (f) CPu (h) NAc. (i) Grouped cilia length (µm) by number of cells (%) in the PFC, CA1, CPu, and NAc.

DREADD hM3Dq-mediated chemogenic activation of MCH neurons in pMchCre+ adult mice caused a significant decrease in cilia length in comparison to pMchCre- adult mice in both hemispheres of the brain in several regions of the brain including the CA1, striatum, PFC, and NAc. Co-expression of mCherry fluorescence (red) identifying DREADD-expressing neurons and MCH immunofluorescence (green) in PmchCre+ and PmchCre- mice was confirmed (Fig. 4b). Coexpression of mCherry fluorescence (red) identifies DREADD-expressing neurons and c- Fos immunofluorecense (green) confirmed cells recently activated in PmchCre+ and PmchCre- mice (Fig. 4c). In the PFC stimulating MCH neurons in pMchCre- mice caused a 40% decrease in cilia length (5.226 ± 0.4111 µm in pMchCre- mice compared to 3.125 ± 0.3080 µm in pMchCre+ mice, t=4.090, P=0.0064, unpaired t-test, Fig 4d,e). In the CA1 stimulating MCH neurons in pMchCre- mice caused a 39% decrease in cilia length (6.079 ± 0.3029 µm in pMchCre- mice compared to 3.699 ± 0.3173 µm in pMchCre+ mice, t=5.426, P=0.0016, unpaired t-test, Fig 4f,g). In the CPu stimulating MCH neurons in pMchCre- mice caused a 40% decrease in cilia length (6.792 ± 0.3493 µm in pMchCre- mice compared to 4.059±0.4322 µm in pMchCre+ mice (t=4.918, P=0.0027, unpaired t-test, Fig 4h,i). In the NAc stimulating MCH neurons in pMchCre- mice caused a 32% decrease in cilia length (9.240±0.4591 µm in pMchCre- mice compared to 6.267±0.1699 µm in pMchCre+ mice, t=6.073, P=0.0009, unpaired t-test, Fig 4j,k). Number of cells (%) were grouped by cilia length (µm) in the PFC, CA1, CPu, and NAc to further show the decrease in cilia length when MCH is stimulated via chemogenics (Fig. 4l). Stimulating MCH neurons via chemogenics caused a significant decrease in cilia density per field in the PFC (t=4.092, P=0.0064) and CA1 (t=3.613, P=0.0112, Fig. S4).

### 3. Inactivation of the MCH system increases cilia length *in the mouse brain*

#### 3.1. MCHR1 antagonist lengthens cilia

The administration of GW803430 via intraperitonial injection in adult mice for 7 consecutive days caused a significant increase in the cilia length in multiple brain regions including the CA1 of the CA1, striatum, PFC, and NAc (Fig 5a). In the PFC, administration of GW803430, caused cilia an increase in cilia length by 42% (6.845 ± 0.3 µm compared to 3.935 ± 0.42 µm in the PFC of the animals administered vehicle, t=5.646, P=0.0013, unpaired t-test, Fig 5b,c). In the CA1 administration of GW803430 caused an increase in cilia length by 28% (8.595 ± 0.55 µm compared to 6.103 ± 0.5 µm in the CA1 of the animals administered vehicle, t=3.4, P=0.0145, unpaired t-test, Fig 5d,e). In the CPu administration of GW803430 caused lengthening in cilia by 33%. (7.30 ± 0.50 µm compared to 4.84 ± 0.24 µm in the CPu of the animals administered vehicle, t=4.583, P=0.0038, unpaired t-test, Fig 5f,g). In the NAc administration of GW803430 caused an increase in cilia length by 17% (9.190 ± 0.17 µm compared to 7.544 ± 0.22 µm in the NAc of the animals administered vehicle, t=5.992, P=0.0010, unpaired t-test, Fig 5h,i). Number of cells (%) were grouped by cilia length (µ) in the PFC, CA1, CPu, and NAc to further show the increase in cilia length when GW803430 is administered i.p. in mice (Fig. 5j). Administration of GW resulted in no significant difference in cilia density per field in the PFC, CA1, CPu, and NAc (Fig, S5).

**Fig.5.**
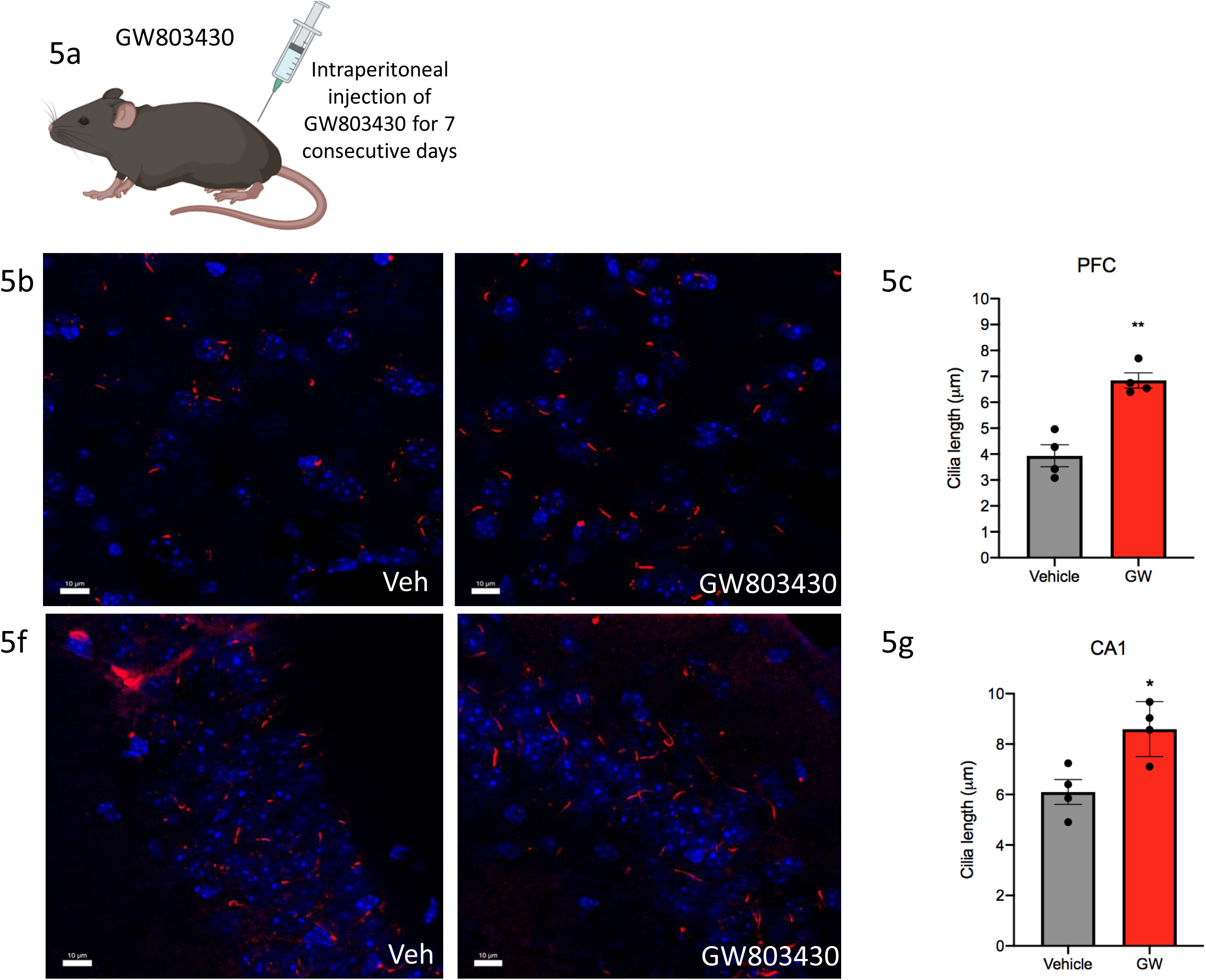
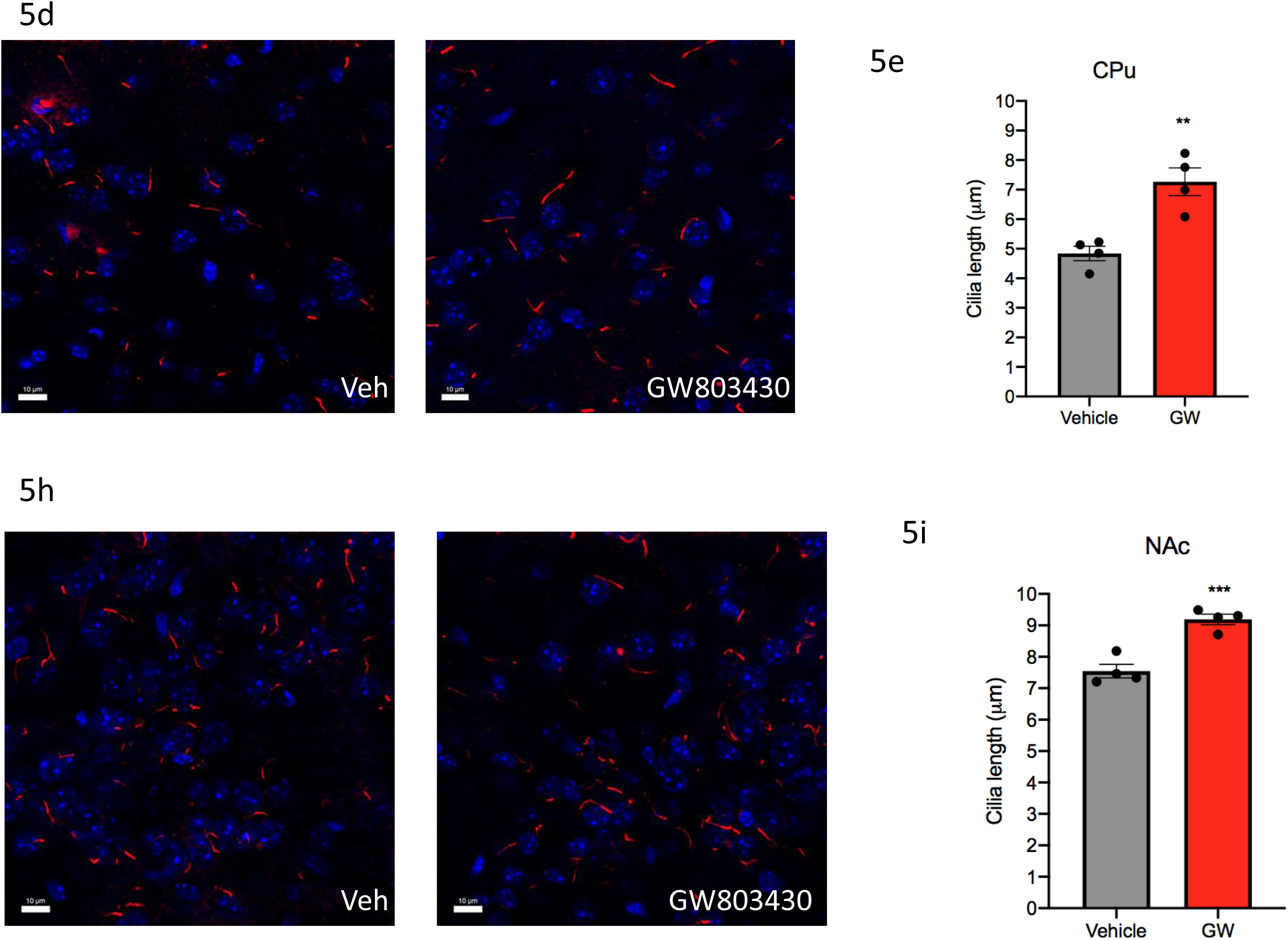
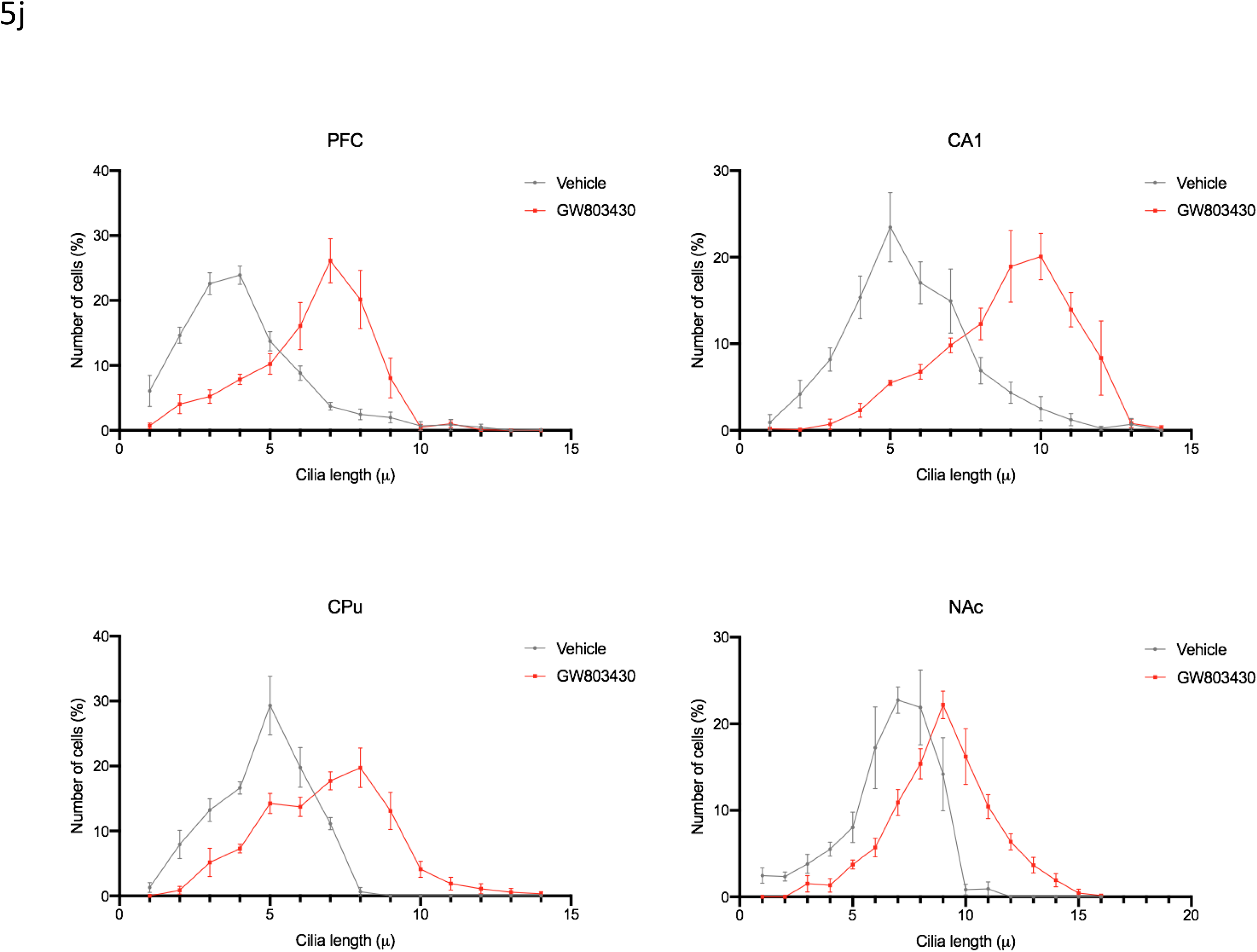
Cilia length is increased in MCH deficit mice. (a) Schematic representation of how the MCH cells are ablated through DTR. IDTR mice were crossed with PmchCre mice rendering offspring mice with Cre expressing MCH neurons sensitive to DT. Created with BioRender.com (b) MCH immunoreactivity (GFP) in the lateral hypothalamus of IDTRPmchCre- and IDTRPmchCre+ mice following DT injection (Scale bar= 100 µm) Immunostaining of ADCY3 labeled in red fluorescence in IDTRPmch+ and IDTRPmch- mice in the (c) PFC, (e) CA1, (g) CPu and (i) NAc (Scale bar= 10 µm). Quantification of cilia length (µm) in the (d) PFC, (f) CA1, (h) CPu (j) NAc. (k) Grouped cilia length (µm) by number of cells (%) in the PFC, CA1, CPu, and NAc.

#### 3.2. Cilia length is increased in MCH and MCHR1 deficit mice

Germline deletion of MCHR1 in mice caused a significant increase in the cilia length in several regions of the brain including CA1, CPu, and PFC (PFC) and NAc. In the PFC of MCHR1 KO mice there was an increase in cilia length by 41% (6.50±0.09 µm in the MCHR1KO compared to 3.8±0.1 µm in WT mice, t=20.15, *P* <0.0001, unpaired t-test, Fig 6a,b). In the CA1 of the hippocampus of MCHR1 KO mice there was an increase in cilia length by 22% (3.6±0.10 µm. 0.1 µ compared to 2.8± 0.03 µm in the CA1 of the WT mice (t= 6.712, *P* = 0.0003, unpaired t-test, Fig 6c,d). In the CPu of the MCHR1 KO mice, cilia length increased by 17% (9.5± 0.3 µm compared to 7.9±0.3 µm in WT mice, t= 3.952, *P* = 0.005, unpaired t-test, Fig 6e,f). In the NAc of the MCHR1 KO mice, cilia length increased by 25% (10.19± 0.50 µ in MCHR1KO compared to 7.60± 0.50 µm in the NAc of the WT mice, t=4.434, *P* = 0.003, unpaired t-test) (Fig 6g,h). Number of cells (%) were grouped by cilia length (µm) in the PFC, CA1, CPu, and NAc to further show the increase in cilia length in MCHR1 KO mice (Fig. 6i). MCHR1KO mice resulted in a significant increase in cilia density per field in the CA1 (t=3.884, P=0.0081, Fig. S6).

**Fig.6.**
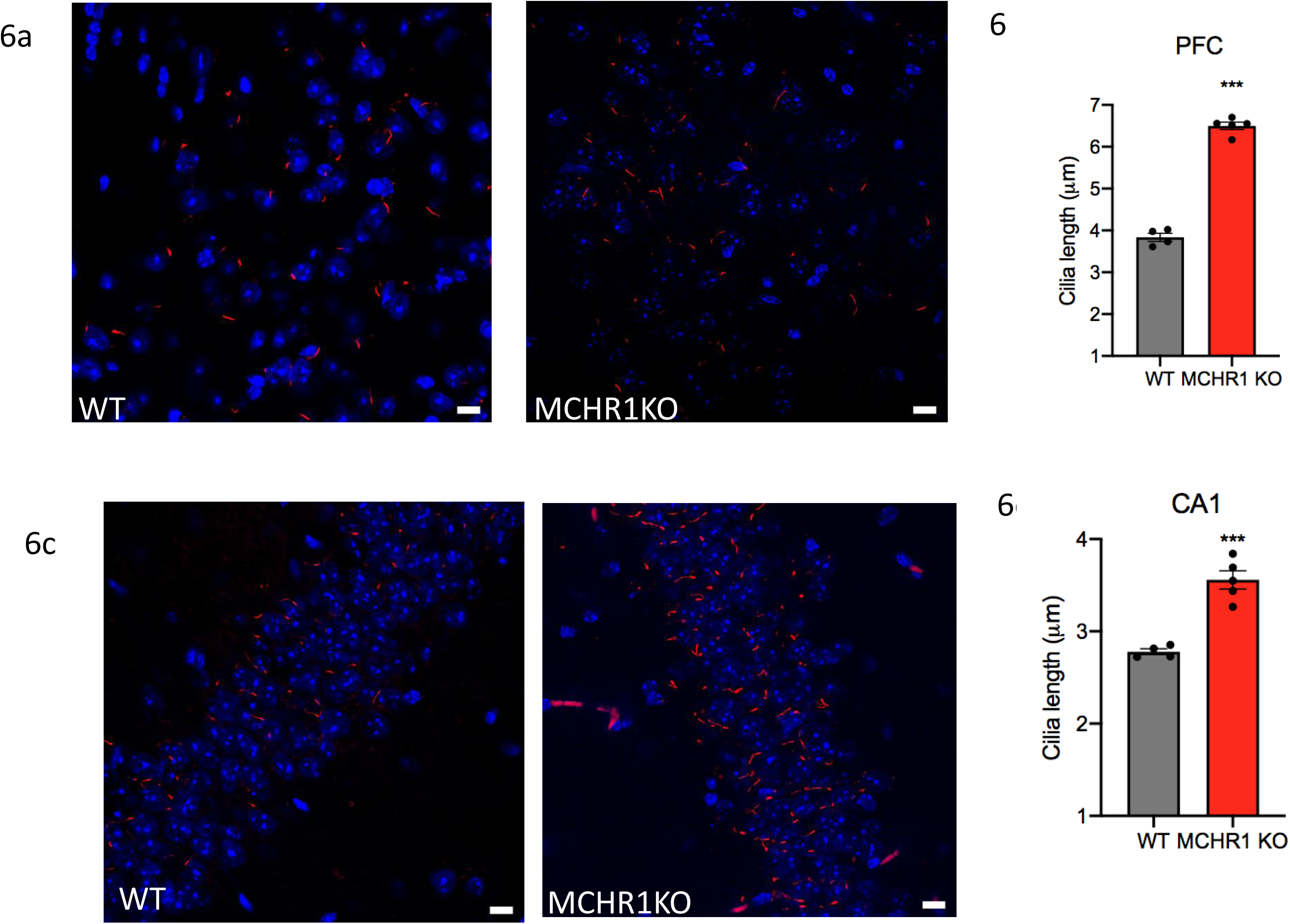
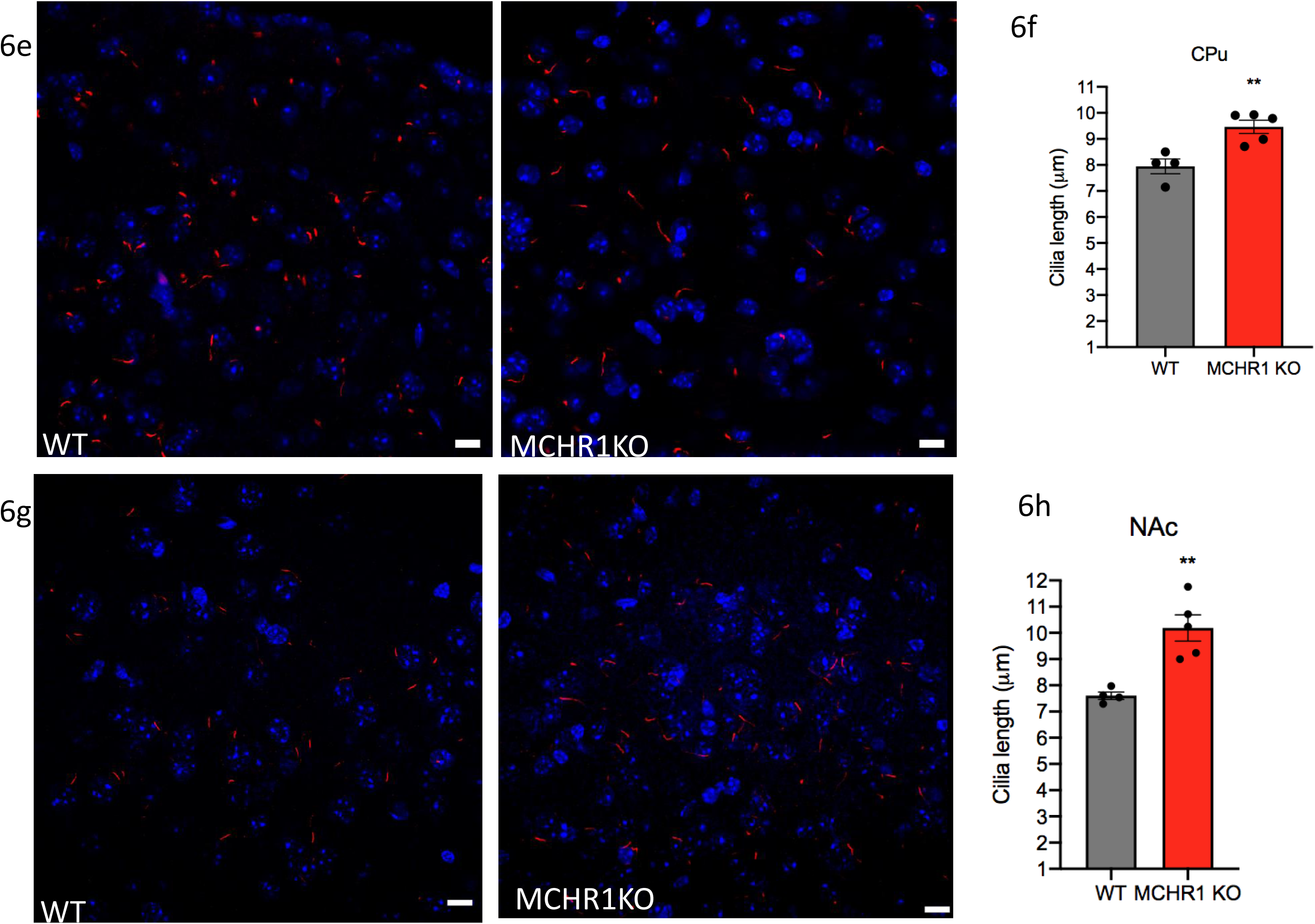
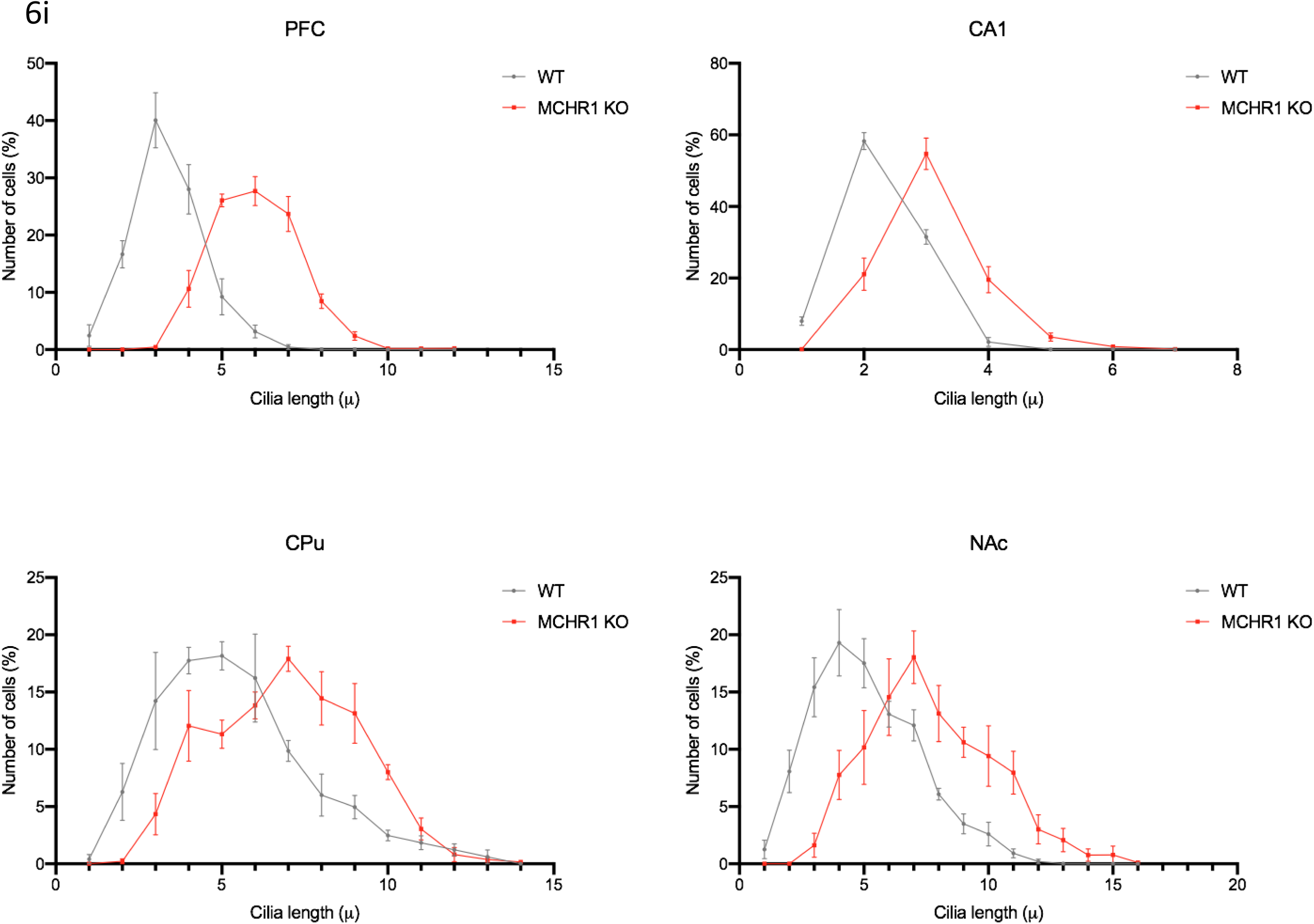
Optogenetic stimulation of MCH shortens cilia length. (a) Experimental approach. Adult PmchCre+ and PmchCre- mice were stereotaxically injected with AAV-EF1α-DIO-ChR2-T159c-eYFP in the lateral hypothalamus (LH). A fiber cannula was then placed in the LH slightly above the injection site. Created with BioRender.com (b) Coexpression of ChR2 (green) containing neurons in the lateral hypothalamus and MCH immunofluorescence (red) in PmchCre+ and PmchCre- mice. (Scale bar= 10 µm). (c) ChR2 fluorescence (green) and c-Fos immunofluorecense (red) identifies cells recently activated in PmchCre+ and PmchCre- mice (Scale bar= 10 µm). Immunostaining of ADCY3 labeled in red fluorescence in the (d) PFC, (f) CA1, (h) CPu and (j) NAc via CNO-dependent activation of Gq signaling in PmchCre+ and PmchCre- mice (Scale bar= 10 µm). Quantification of cilia length (µm) in the (e) PFC, (g) CA1, (i) CPu (k) NAc. (l) Grouped cilia length (µm) by number of cells (%) in the PFC, CA1, CPu, and NAc.

The conditional ablation of MCH neurons in adult mice (MCH-cKO) caused a significant increase in the cilia length in several regions of the brain including CA1, striatum, and PFC and NAc (Fig 7a,b). In the PFC of MCH-cKO mice, cilia length increased by 19% (6.185 ± 0.04609 µm in IDTR+pMchCre+ mice compared to 5.03 ± 0.32 µm in IDTR+pMchCre- mice, t=3.539, P = 0.024, unpaired t-test, Fig 7c,d). In the CA1 of IDTR+MCH-cKO mice, cilia length increased by 24% (8.209 ± 0.07566 µm in IDTR+pMchCre+ mice compared 6.24 ± 0.23 µm in IDTR+pMchCre- mice, t=8.265, P = 0.0012, unpaired t-test) (Fig 7e,f). In the CPu of MCH-cKO mice, cilia length increased by 29% (7.373 ± 0.6131 µm in IDTR+pMchCre+ mice compared to 5.242 ± 0.2518 µm IDTR+pMchCre- mice, (t=3.215, *P* = 0.0324, unpaired t-test, Fig 7g,h). In the NAc of MCH-cKO mice, cilia length increased by 23% (6.823± 0.3695 µm in IDTR+pMchCre+ mice compared to 5.239± 0.4177 µm in IDTR+pMchCre- mice (t=2.841, *P* = 0.0456, unpaired t-test) (Fig 7i,j). Number of cells (%) were grouped by cilia length (µm) in the PFC, CA1, CPu, and NAc to further show the increase in cilia length in MCH-cKO mice (Fig. 7k). MCHcKO mice resulted in a significant increase in cilia density per field in the CA1 (t=6.516, P=0.0006) and the NAc (t=5.047, P=0.0023, Fig. S7).

**Fig.7.**
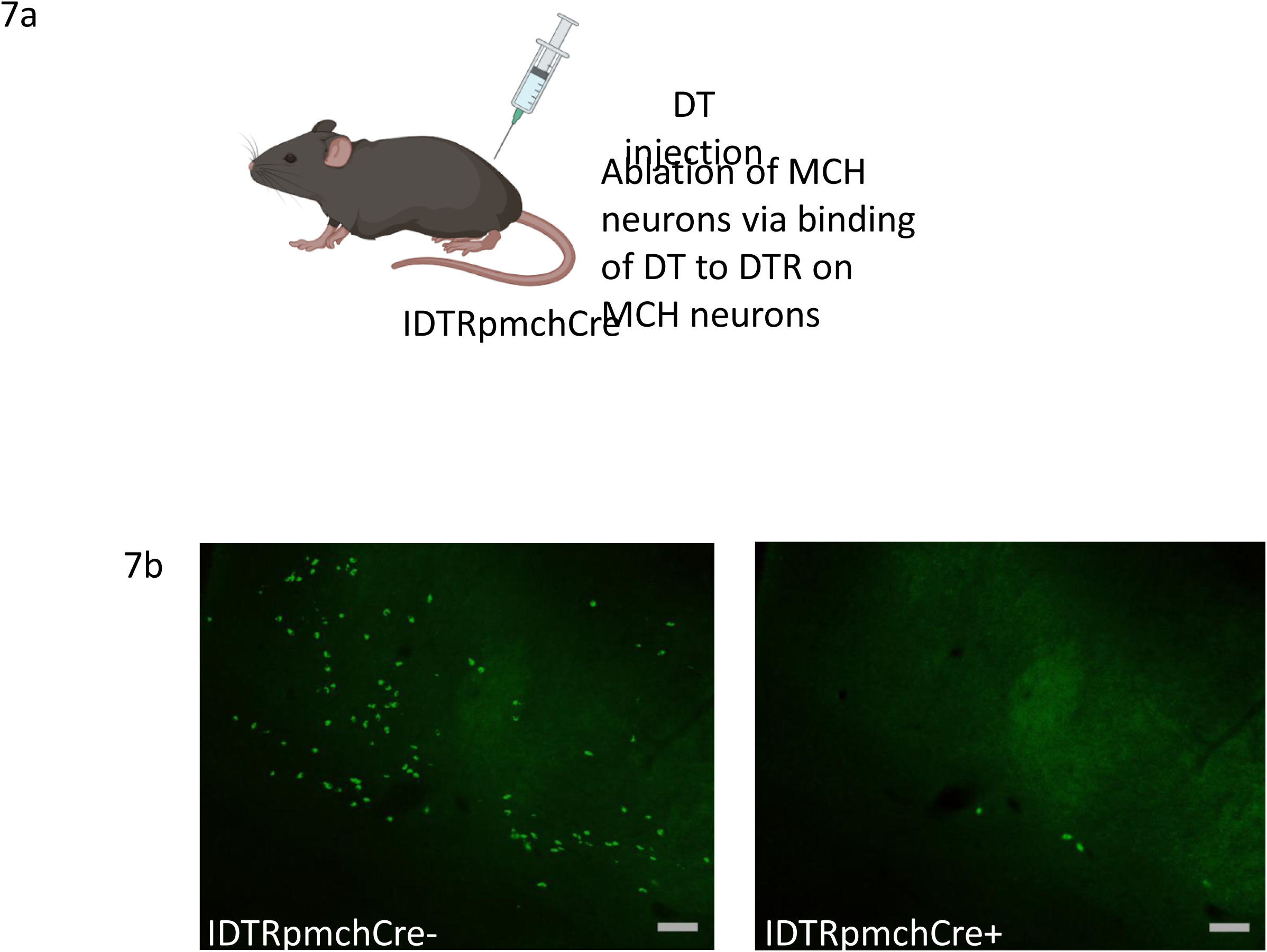
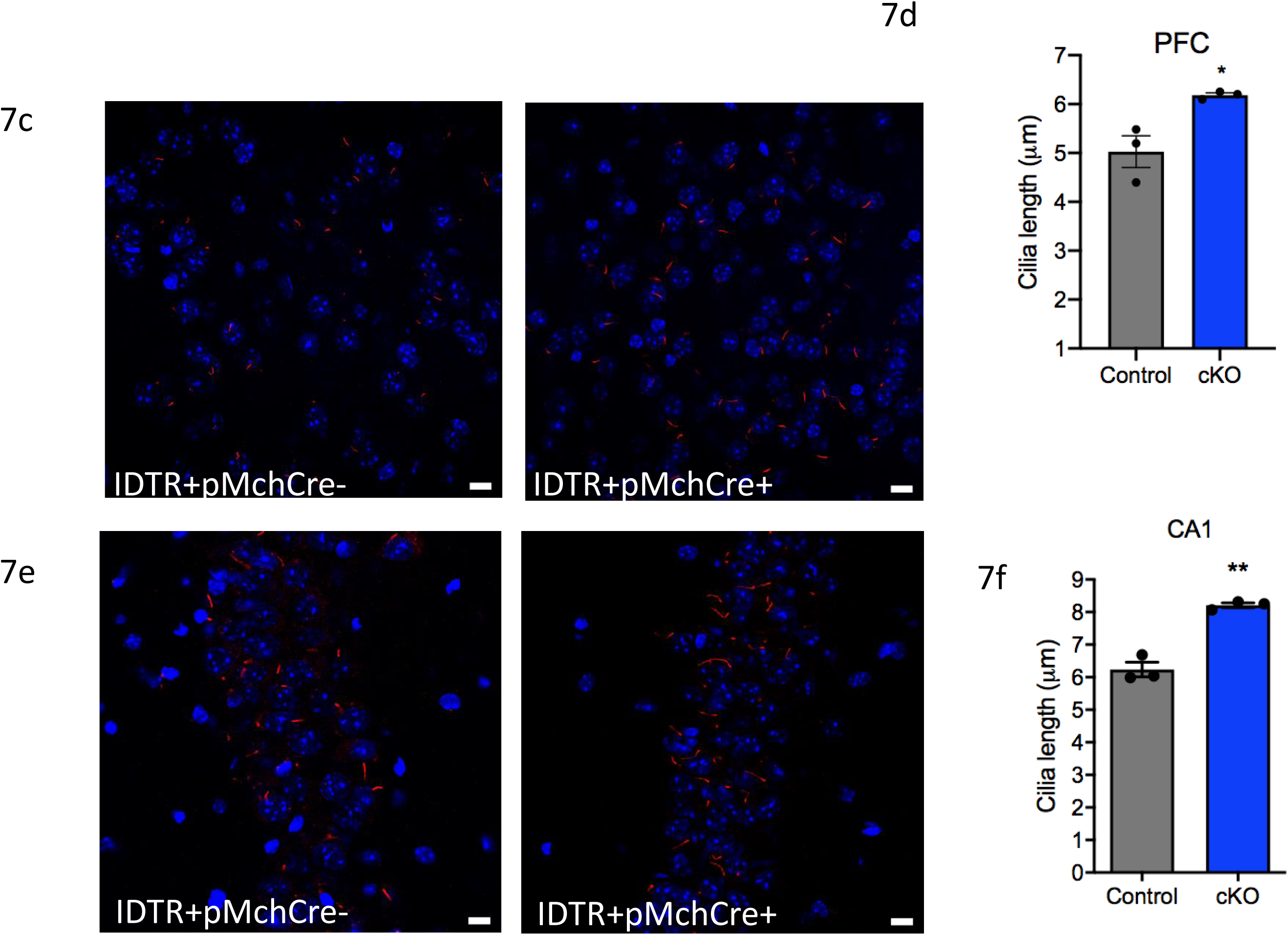
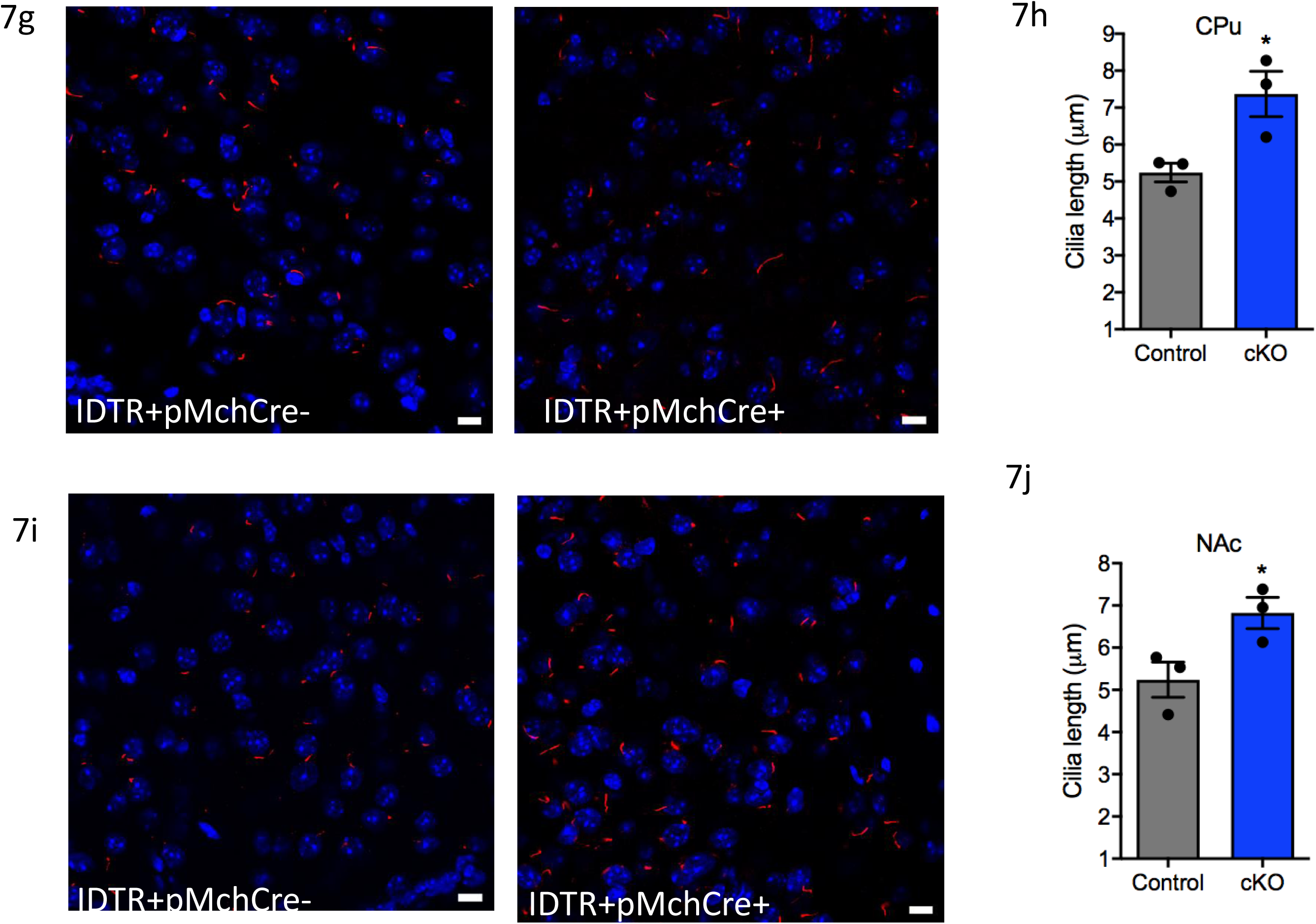
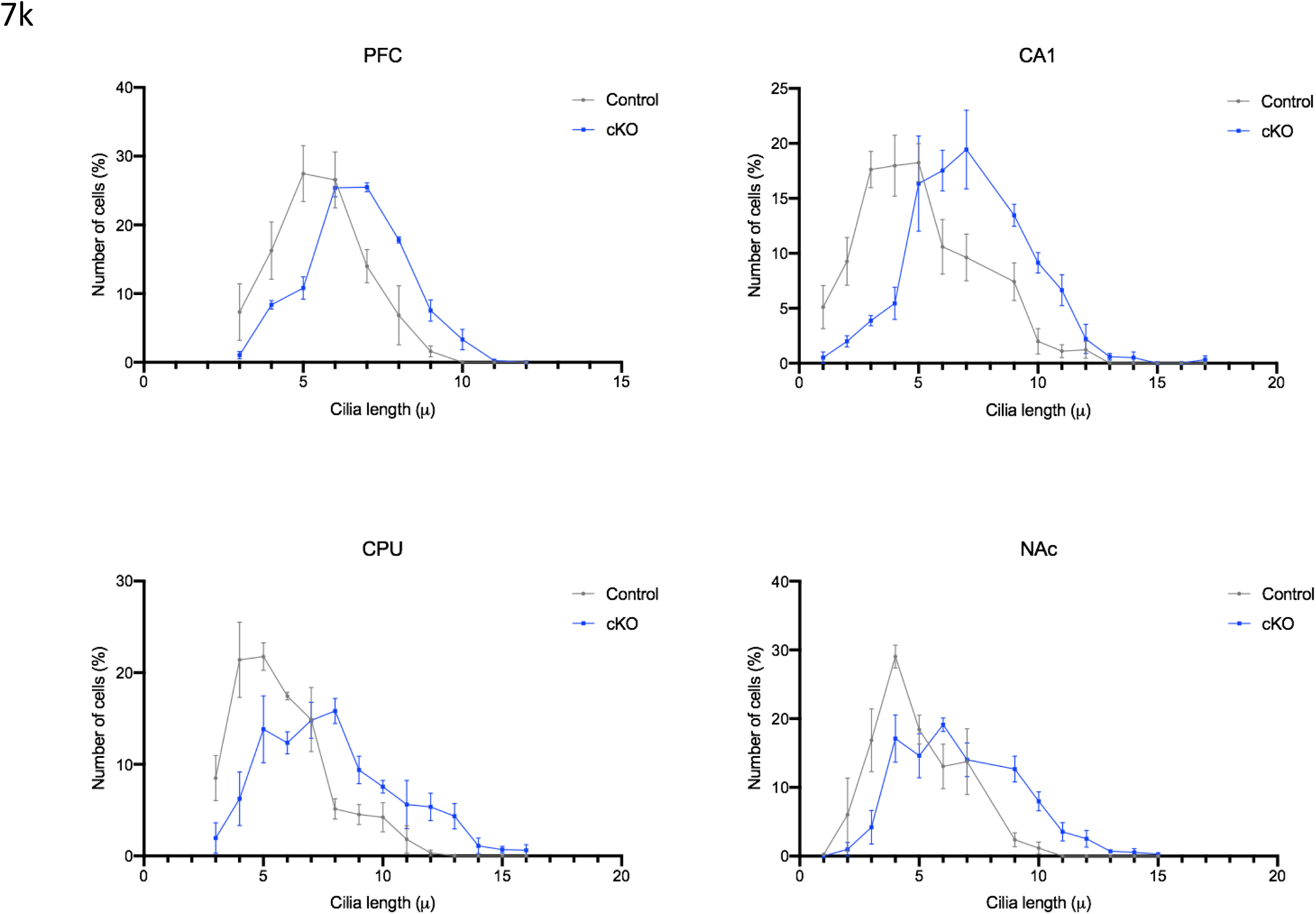
Chemogenic excitation of MCH shortens cilia length. (a) Experimental approach. Adult PmchCre+ and PmchCre- mice underwent bilateral stereotaxic injection in the lateral hypothalamus with AAV-hSyn-DIO-hM3D(Gq)-mCherry to express DREADD in Cre expressing MCH neurons. Clozapine-N-oxide (CNO) was delivered intraperitoneally to stimulate MCH neurons. Created with BioRender.com (b) Co-expression of mCherry fluorescence (red) identifying DREADD-expressing neurons and MCH immunofluorescence (green) in PmchCre+ and pmchcre- mice (Scale bar= 10 µm). (c) mCherry fluorescence (red) identifies DREADD-expressing neurons and c-Fos immunofluorecense (green) identifies cells recently activated in PmchCre+ and PmchCre- mice (Scale bar= 10 µm). Immunostaining of ADCY3 labeled in red fluorescence in the (d) PFC, (f) CA1, (h) CPu and (j) NAc via CNO-dependent activation of Gq signaling in PmchCre+ and PmchCre- mice (Scale bar= 10 µm). Quantification of cilia length (µm) in the (e) PFC, (g) CA1, (i) CPu (k) NAc. (l) Grouped cilia length (µm) by number of cells (%) in the PFC, CA1, CPu, and NAc.

## Discussion

In this study we established the causal effects of MCH signaling pathways on cilia length. We first showed that MCH treatment causes cilia shortening in the striatal and cortical brain slice cultures. We then demonstrated that the stimulation of MCH system, through direct agonist activation of the MCHR1 or via optogenetic and chemogenic excitation of MCH neurons, causes a significant decrease in cilia length. The inactivation of the MCH system, through pharmacological blockade of MCHR1 or genetic manipulation (MCHR1 germline deletion or MCH neurons’ conditional ablation) causes a significant increase in cilia length.

MCHR1 is widely distributed throughout the brain, with high density in the CPu, NAc, CA1 of the CA1, and the PFC [15, 38]. Therefore, we focused on these regions in our examination of the effects of the activation and inactivation of MCH system on cilia length. One of our unexpected findings was that cilia lengths differ among brain regions and among different mouse strains.

The various manipulation approaches allowed us to: 1) recapitulate conditions used previously to substantiate MCH physiological functions; 2) differentiate between the effects of MCH neurons versus MCHR1 manipulation; 3) distinguish the effects of acute versus chronic activation/inactivation of MCH system; and 4) determine the effects of early-life versus adult stage MCH system manipulations.

The role of MCH in regulating a wide variety of physiological functions such as feeding, obesity, reward, and sleep has been established using pharmacological and genetic manipulations [39–43]. For example, central chronic infusions of MCH has long been shown to induce mild obesity in wild type mice [44], whereas the subchronic administration of MCHR1 antagonists and MCHR1/MCH genetic deletion are known to produce anti-obesity effects in mice [26, 27]. The ciliopathy Bardet-Biedl Syndrome (BBS) is characterized with obesity, and mutations affecting the mouse orthologs of BBS-associated genes disrupt the localization of MCHR1 [45, 46] Thus, failure of MCHR1 to reach the cilium could potentially associate with failure in MCHR1 signaling pathway in the cilia, leading to obesity in BBS [47, 48].

Given that MCH neurons synthesize and release other neurotransmitters/neuropeptides including GABA and NEI and NGE [49, 50], it is important to distinguish the selective role of MCHR1 from that of MCH neurons in regulating cilia length. Hence, MCHR1 pharmacological activation and blockade by MCH central infusion and GW803430 treatment respectively and MCHR1 germline deletion allowed for exploring the selective role of the MCHR1. In parallel, optogenetic and DREADD stimulation of the MCH neurons and IDTR- dependent ablation of MCH neurons allowed for exploring the role of MCH neurons. Optogenetics and DREADD technologies have also been previously used to explore the role of MCH neurons in sleep, reward, and feeding [51–53].

Previous reports demonstrated distinctive effects of acute versus chronic activations of MCH systems on body weight [54]. Therefore we examined the effects of acute (optogenetic and chemogenetic stimulation) and chronic activations (MCH icv administration for 7 days) of the MCH system on cilia length. We previously showed that developmental and adulthood inactivation of the MCH system in mice leads to distinct behavioral deficits. For example, germline MCHR1 deletion but not adult IDTR- dependent ablation of MCH neurons caused olfactory impairment and social deficits [13]. Therefore, we examined here the effects of germline MCHR1 deletion and adulthood conditional MCH ablation on cilia length.

Strikingly, we found that the different MCH system manipulations produced consistent patterns of alterations in cilia length. The activation of MCH system consistently shortened cilia length, while the inactivation of MCH system lengthened cilia length. The findings that both acute and chronic activation of MCH system shortened cilia length and that both early life and adulthood inactivation of MCH system lengthened cilia substantiate the dynamic nature of the cilia system and suggest that cilia’s morphology undergoes rapid changes in response to its environment. Most importantly, our findings provide the first evidence for the direct and moment-to-moment regulation of the brain cilia structure by the MCH system.

Unexpectedly, unilateral optogenetic stimulation of MCH neurons in the lateral hypothalamus caused bilateral shortening of cilia length in the PFC, CA1, CPu, and NAc. This finding may suggest that MCH neurons project ipsilaterally and contralaterally to these brain regions. Another explanation might be that MCH activation may produce ipsilateral modifications of other neurotransmitter systems, which may result in contralateral shortening of cilia length. It is well established that MCH neurons innervate serotonergic neurons in the raphe nucleus and regulate the activity of these neurons [55, 56]. Indeed, specific MCH-regulated functions are mediated through serotonin system [57, 58]. Thus, the activation of MCH neurons may regulate cilia indirectly through its interaction with serotonin system. Interestingly, MCHR1 and 5-HT6 are among the very few GPCRs that preferentially localize to primary cilia [59], and the inhibition of 5-HT6 receptors is known to shorten cilia length [39, 60].

Defects in the assembly such as cilia shortening have been shown to cause a range of severe diseases and developmental disorders called ciliopathies, which are associated with neurological deficits such as abnormal cortical formation and cognitive deficits [61–63]. We have recently shown that that brain cilia genes were differentially expressed in major psychiatric disorders including schizophrenia, and autism spectrum disorder, depression, and bipolar disorder [23]. On the other hand, evidences from our own work and others strongly support an essential role for MCH system particularly MCHR1 in the pathophysiology of schizophrenia. We showed that MCHR1 mRNA is decreased in the PFC of patients with schizophrenia [13]. We also demonstrated that genetic manipulation of the MCH system via the deletion of MCHR1 and the conditional ablation of MCH neurons resulted in behavioral abnormalities mimicking schizophrenia-like phenotypes including repetitive behavior, social impairment, impaired sensorimotor gating, and disrupted cognitive functions [64]. We also showed that MCHR1 germline deletion causes in mice alterations in depressive-like behavior that is sex-dependent [13]. These same genetic models were used in our current study to examine the consequences of these genetic manipulation on cilia length. In both IDTR+pMchCre+ and MCHR1KO animals we found a significant increase in cilia in all regions including the PFC, CA1, CPu, and NAc. Our previous and current data may point at the role of ciliary MCHR1 in the pathopgysiology of psychiatric disorders such as schizophrenia and depression. Together, our previous and current studies point at the cilia elongation as a possible mechanism, through which MCH system dysregulation causes social and cognitive deficits in mice, and is associated with psychiatric disorders in humans [13, 23]. Our results also prompt the question on whether selective targeting of cilia MCHR1 might offer a novel approach to treat psychiatric disorders. Indeed, studies have demonstrated that some pharmacological agents can alter cilia length. For example, lithium, which is a mood stabilizer that is used in treatment of bipolar disorder and acute mania, increases MCHR1-positive cilia length in several cell types including neuronal cells, specifically the dorsal striatum and NAc [65].

In conclusion, our present study has established the causal regulatory effects of the MCH system signaling on cilia length. The findings of this study are significant because (1) they show for the first time in vivo that primary cilia length can be regulated by the MCH signaling, proving the link between GPCRs and cilia length regulation, and (2) they implicate ciliary MCHR1 as a potential therapeutic target for the treatment of pathological conditions characterized by impaired cilia function.

## Supplemental Figures

**Fig. S1** Stimulation of MCH increases food intake via optogenetics. PMchCre+ mice spend more time eating than pMchCre- mice. Over the observed 10 minutes, pMchCre+ mice averaged 74.13 seconds of feeding while pMchCre- mice averaged 7.413 seconds (t=7.027, P<0.001)

**Fig. S2** Administration of MCH i.c.v. results in a decrease in cilia density per field in PFC (t=3.849, P=0.0085), CA1(t=3.141, P=0.02), CPu (t=4.224, P=0.0055), and NAc (t=3.766, P=0.0093)

**Fig. S3** Stimulating MCH neurons via optogenetics caused a decrease in cilia density per field in PFC (t=3.789, P=0.0091), CA1 (t=6.324, P=0.0007), and CPu (t=5.656, P=0.0013)

**Fig. S4** Stimulating MCH neurons via chemogenics caused a decrease in cilia density per field in PFC (t=4.092, P=0.0064) and CA1 (t=3.613, P=0.0112)

**Fig. S5** Administration of GW resulted in no significant difference in cilia density per field in PFC, CA1, CPu, and NAc

**Fig. S6** MCHR1 KO mice have an increase in cilia density per field in CA1 (t=3.884, P=0.0081)

**Fig. S7** MCHcKO mice have an increase in cilia density per field in CA1 (t=6.516, P=0.0006) and NAc (t=5.047, P=0.0023)

