## Supplementary figures and images for "Regulation of Brain Primary Cilia Length by MCH Signaling: Evidence from Pharmacological, Genetic, Optogenetic and Chemogenic Manipulations"

### Supplemental Figures

#### Supplemental Figures

S1

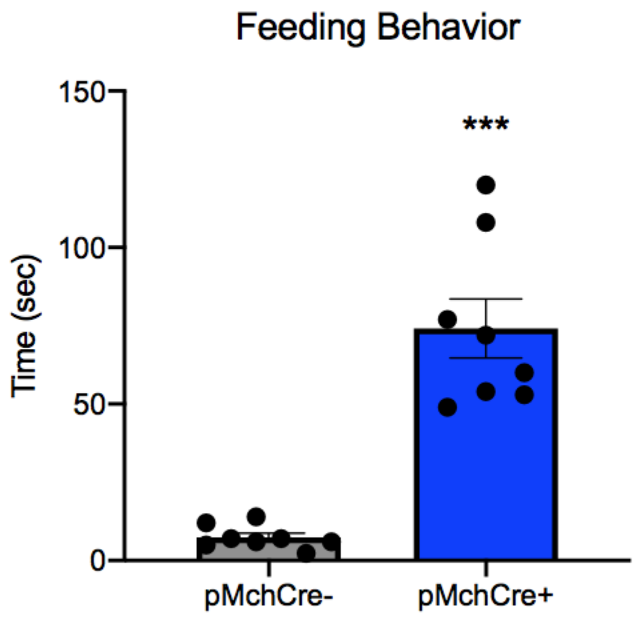

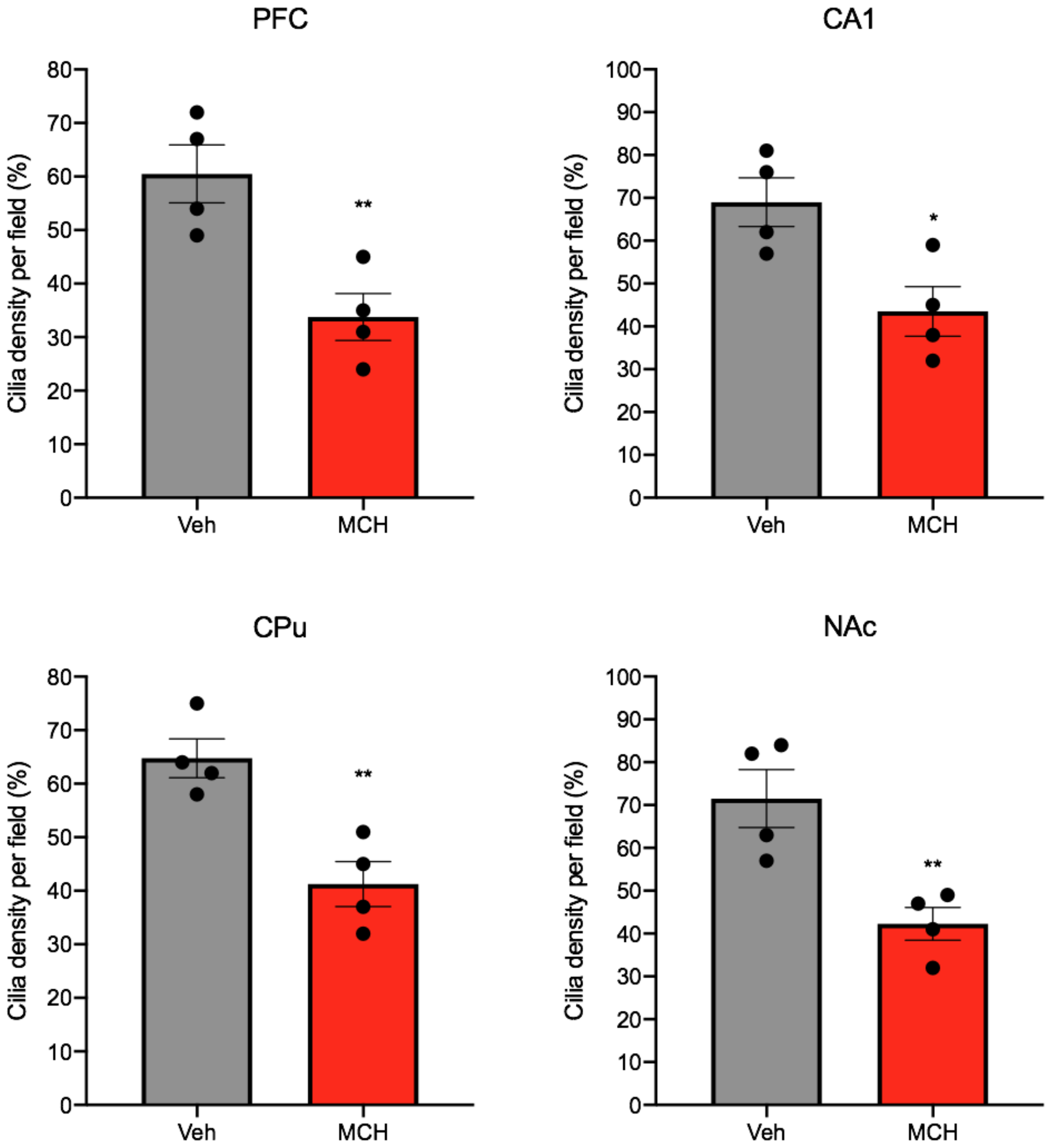

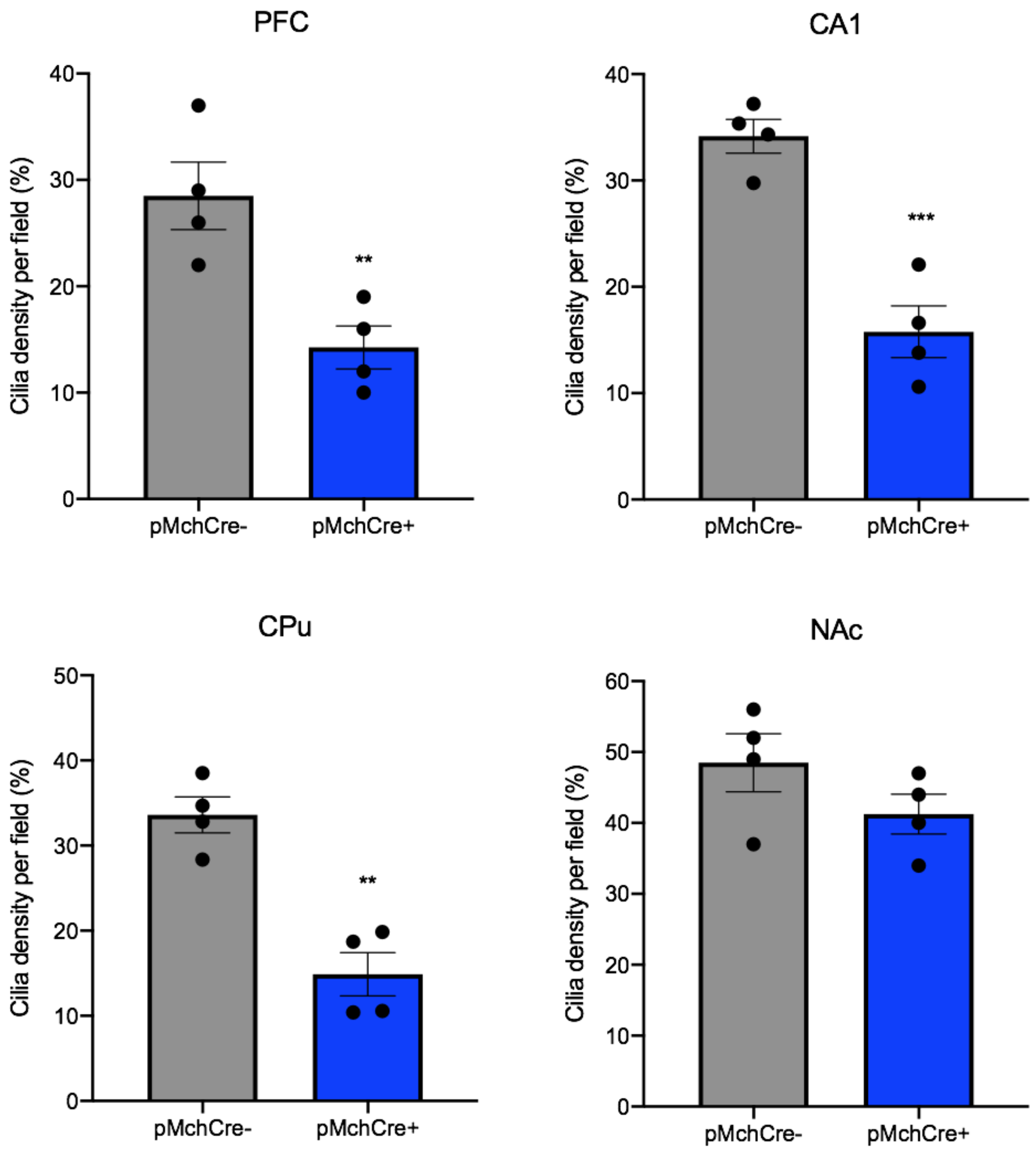

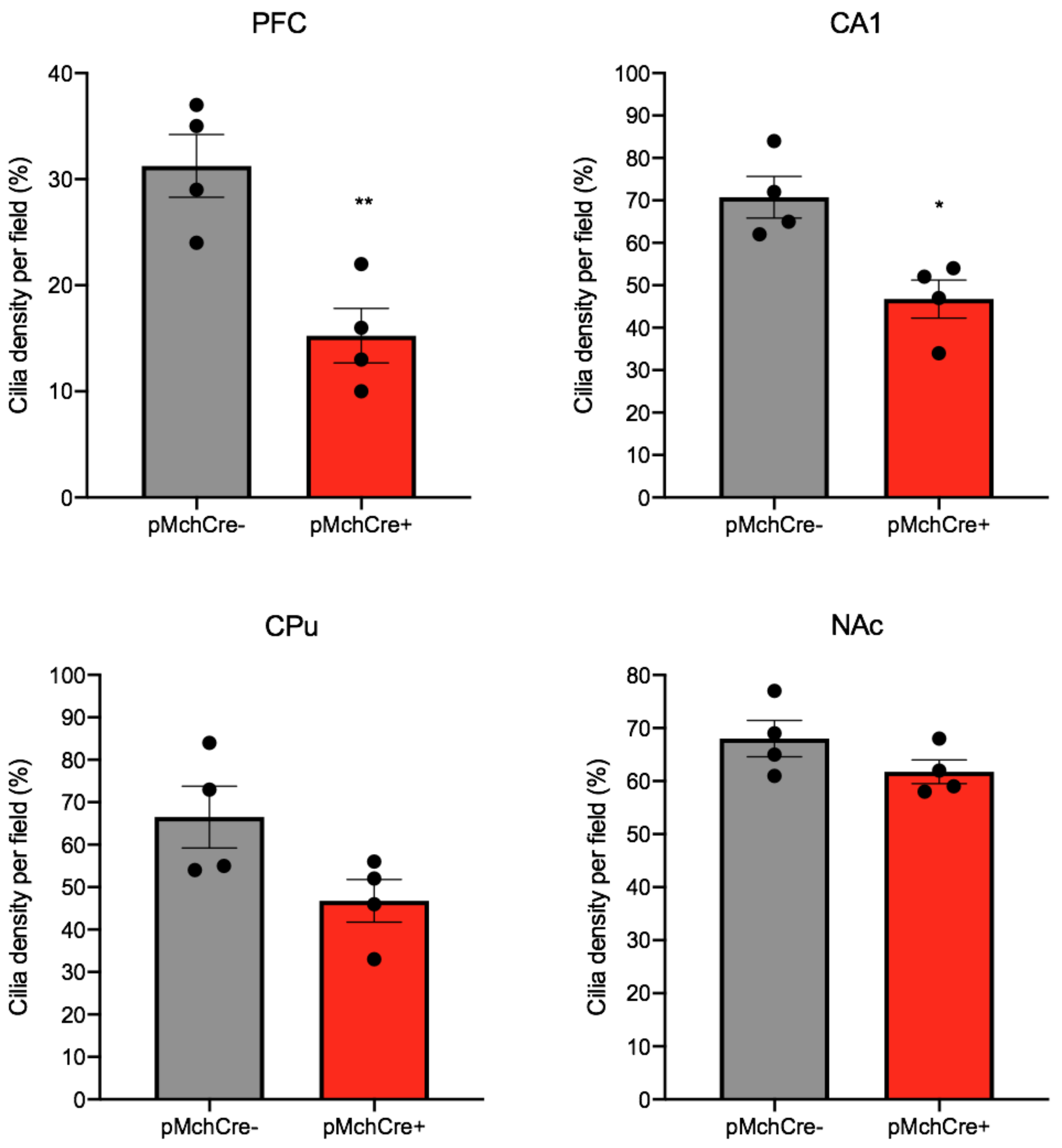

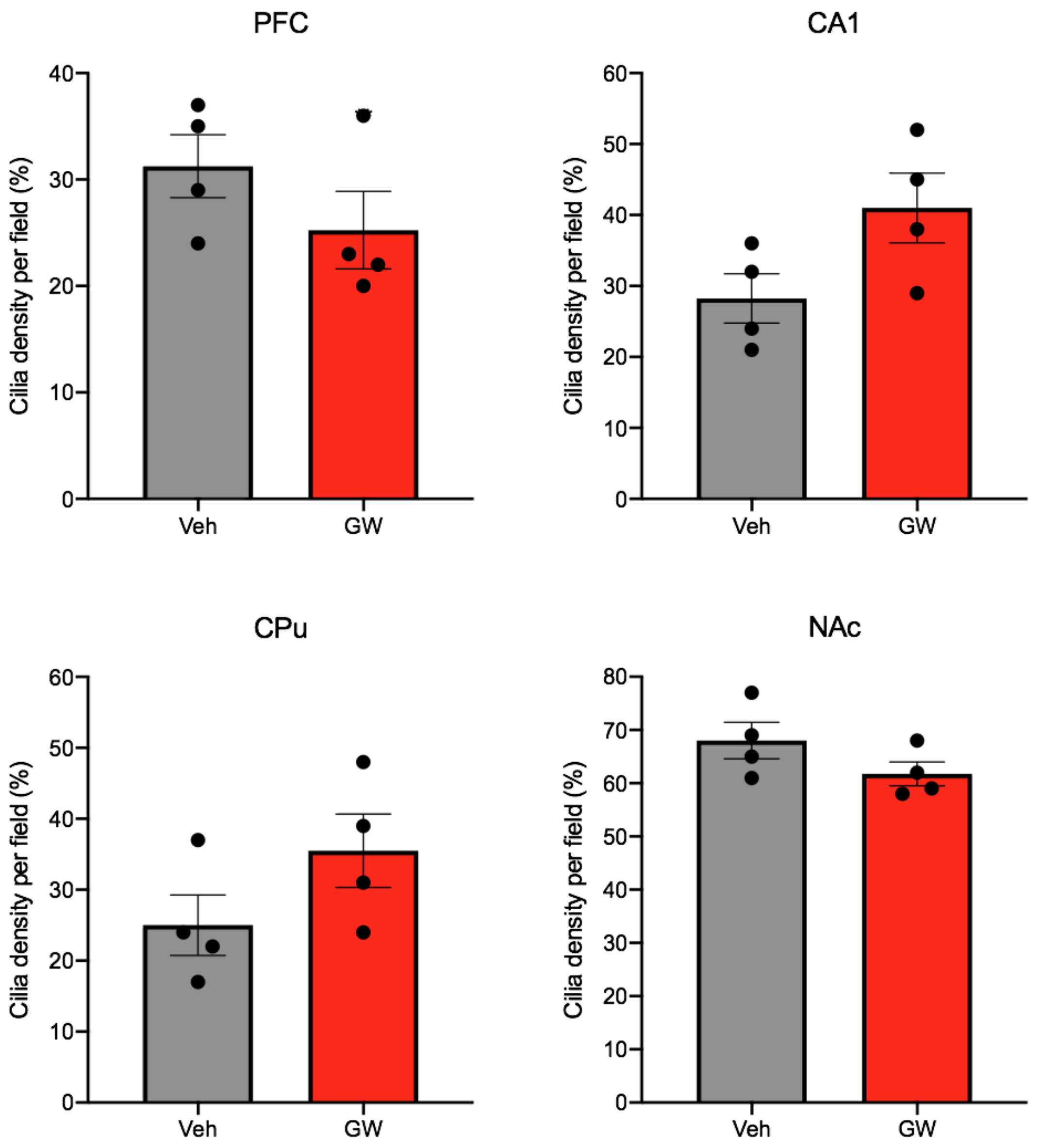

S6

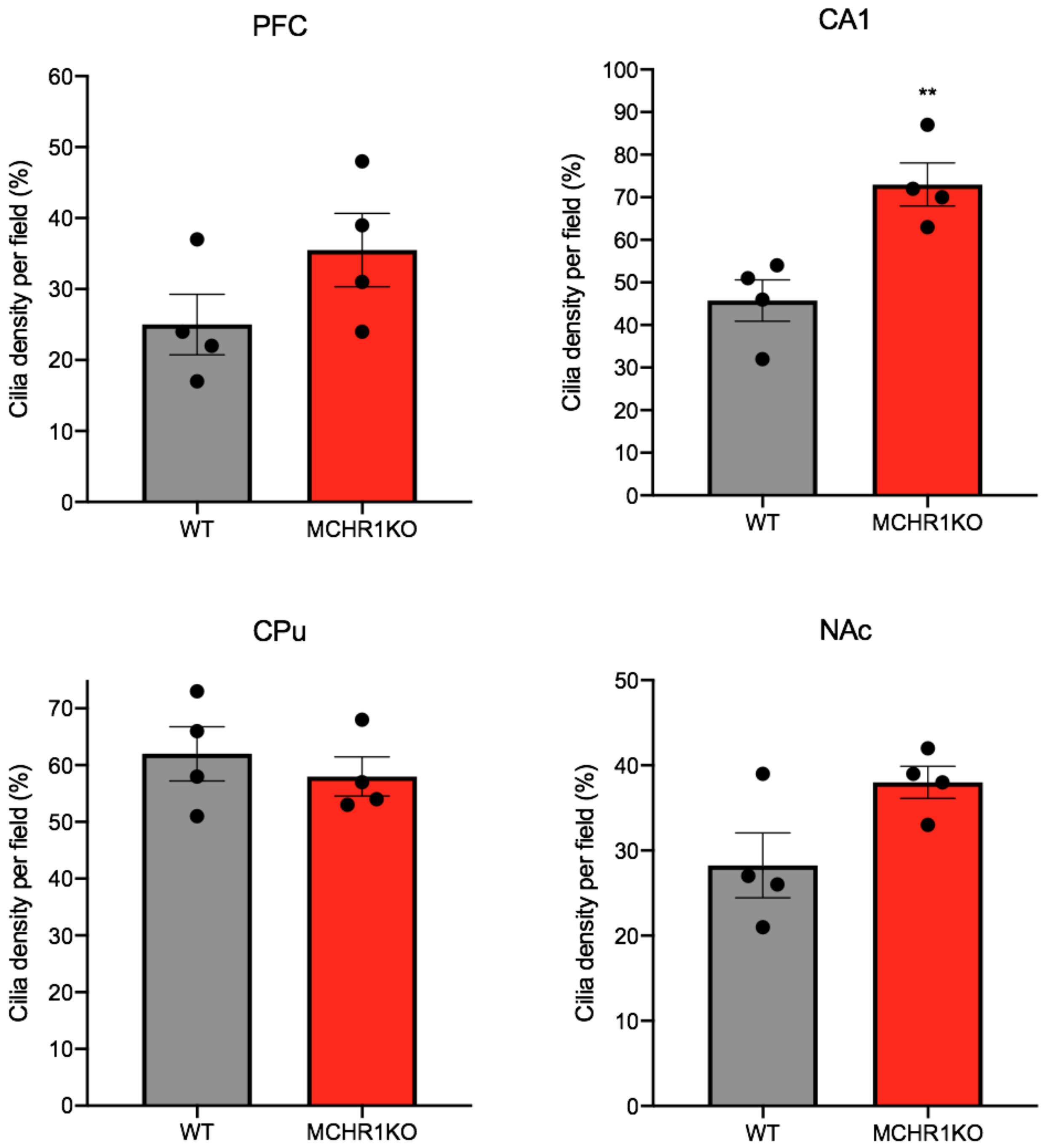

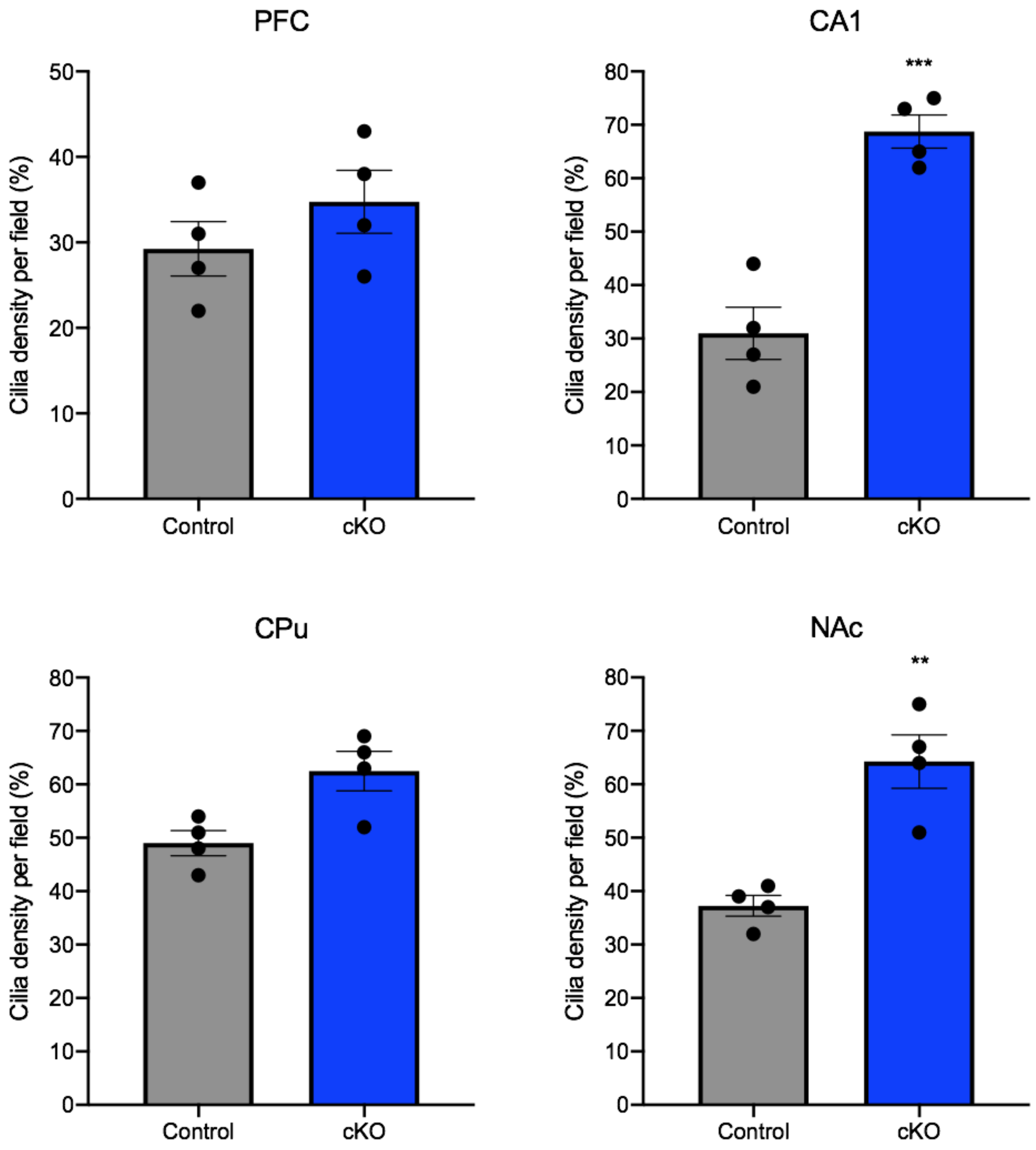
